# Aberrant cytoplasmic localization of MLH1 characterizes a sub-clonal breast cancer cell population that seeds recurrence

**DOI:** 10.1101/2024.02.27.582389

**Authors:** A Mazumder, JT DeWitt, E Oropeza, N Punturi, D Lozano, M Raghunathan, JM Piscitelli, E Sajjadi, E Guerini-Rocco, K Venetis, M Ivanova, E Mane, N Fusco, MN Bainbridge, CM Manhart, S Haricharan

## Abstract

Estrogen receptor positive (ER+) breast cancer is one of the most common causes of cancer-related death in women. Mortality is largely driven by recurrence of treatment-resistant disease after many years of apparent response, making the molecular events that cause recurrence a critical area of investigation. Loss of expression of *MLH1*, a tumor suppressor best studied in its role in mismatch repair, induces resistance of ER+ breast cancer cells to standard estrogen-targeting therapies. It does so by delinking cell cycle progression from estrogen regulation, a role distinct from its function in mismatch repair. MLH1 loss, as currently clinically diagnosed by detecting genomic instability or by immunohistochemistry for absence of protein, occurs in 12-15% of all cancers. Here, we demonstrate that sub-clonal, patient-derived mutations in *MLH1*, which neither impact protein abundance nor contribute sufficiently to genomic instability to be detected diagnostically, seed endocrine treatment resistance by enabling estrogen-independent growth *in vitro*, *ex vivo* in patient-derived organoids (p=0.005) and *in vivo* (p=0.0001). The mechanism underlying this endocrine treatment resistance is aberrant localization of MLH1 to the cytoplasm *in vitro* and *in vivo* (p=0.04), which precludes cell cycle arrest in response to endocrine therapy while simultaneously rendering cells acutely dependent on CDK4/6 activity. Consequently, administration of CDK4/6 inhibitors causes extreme regression in cells with cytoplasmic MLH1 compared to control cell populations with nuclear localization of MLH1 *in vitro* (p=0.00000009), *ex vivo* (p=0.01) and *in vivo* (p=0.01). As aberrant cytoplasmic localization occurs in an additional ∼12% of ER+ breast cancer patients, it constitutes a new, major contributor to MLH1 dysregulation. The potential applicability of cytoplasmic MLH1 as a predictor of responsiveness to existing targeted therapies in a hard-to-treat breast cancer subtype posits an update of current clinical diagnostic criteria and therapeutic strategies. This is particularly important in the adjuvant setting where identification of biomarkers predicting responsiveness to CDK4/6 inhibitors remains an urgent, unmet clinical need.

## Introduction

Almost 25% of Estrogen Receptor positive (ER+) breast cancer patients, one of the largest cohorts of cancer patients globally, who are initially responsive to standard endocrine-targeting therapies are mismatch repair (MMR) deficient at the time of disease relapse^1, 2^. MMR is a tumor suppressor pathway commonly dysregulated across cancer types principally through the loss of function of one of four principal genes: *MLH1*, *MSH2*, *MSH6*, or *PMS2*^3^. MMR deficient tumors display a molecular phenotype characterized by high numbers of somatic point mutations and genetic instability of numerous microsatellite repeat sequences throughout the genome referred to as microsatellite instability^4^. Microsatellite instability was first described in inherited tumors associated with Lynch syndrome caused by germline mutations in one of the four MMR genes detailed above^5^. MMR loss has independent prognostic value and predicts response to standard therapies across cancer types^6^. Although best studied in the context of Lynch syndrome, MMR loss can also occur somatically in tumors through promoter hypermethylation, mutation or loss of expression of MMR genes^1, 7, 8^. Given its prognostic and predictive value, MMR loss is routinely diagnosed in the clinic either by measuring microsatellite instability and tumor mutation burden, or by immunohistochemistry to measure presence or absence of MMR proteins^3, 9^. However, its utility as a diagnostic differs based on cancer type^10^.

ER+ breast cancer does not fall under the Lynch syndrome spectrum of cancers as it rarely associates with germline mutations in MMR genes^11^. However, somatic loss of gene expression of *MLH1* in ER+ breast cancers induces resistance to endocrine therapy^7, 12, 13^. Since somatic loss of *MLH1* is detectable in ∼30% of endocrine treatment resistant ER+ breast cancer patients, it is an important driver of poor patient outcome^2, 7^. Despite the importance of MLH1 loss as a driver of ER+ breast cancer outcome, little is known about the pathogenicity of somatic mutations in *MLH1.* The majority of *MLH1* mutations in ER+ patient tumors, are missense and non-recurrent making determination of variant pathogenicity burdensome^14–19^. Nevertheless, understanding how *MLH1* mutations impact endocrine treatment resistance could significantly improve the predictive value of identifying these mutations through routine clinical diagnostics, for ER+ breast, and other, cancer patients^18^. Studies that investigate the functional impact of *MLH1* mutations thus far have focused on their role in maintaining mismatch repair efficacy^18, 20^. However, we and others have shown that a key consequence of MLH1 loss in ER+ breast cancer is ATM/Chk2-mediated cell cycle dysregulation rather than decreased mismatch repair^2, 7, 21, 22^. The functional consequence of mutations in *MLH1* on cell cycle regulation and endocrine treatment resistant growth remains virtually unstudied^14, 23^.

Endocrine treatment resistance is a major clinical challenge in treating ER+ breast cancer patients and a primary cause of breast cancer-related death^12^. Sub-clonal diversity is a key contributor to endocrine treatment resistance^24, 25^. Indeed, there is strong evidence in the literature demonstrating that acquired drug resistance in ER+ breast cancer is largely driven by drug-refractory sub-populations (subclones) within these heterogeneous tumors^26–29^. Existence of these subclones contributes to prevalence of low variant allele fraction (VAF) mutations, which are hard to detect routinely without very high coverage of the tumor. Therefore, it is likely that VAF mutations remain undetected for many years, seeding eventual treatment resistance and recurrence. Resistant subclones can be pre-existing (where they remain non-responsive to treatment early on) or acquired (where new sub-clones are created and selected partly due to the strong pressure of treatment)^12, 30–34^. Missense mutations in *MLH1* are generally sub-clonal at diagnosis, and therefore have low VAFs^10, 35^, and since MLH1 loss induces endocrine treatment resistance, they could constitute such persister niches. Understanding their impact on endocrine treatment resistance is a clinically relevant question that could potentially prevent deaths of thousands of ER+ breast cancer patients every year around the world.

About 20% of ER+/HER2− breast cancer patients are intrinsically resistant to endocrine therapy while ∼40% of initially responsive patients acquire resistance over time^36^. For patients who are endocrine treatment resistant, CDK4/6 inhibitors such as palbociclib, ribociclib and abemaciclib have emerged as one of few targeted agents with quantifiable benefit^37^. However, although the combination of CDK4/6 inhibitors and endocrine therapy improves median progression-free survival in metastatic endocrine treatment resistant patients, it does not prolong overall survival^37^. This is likely because many patients who initially benefit from CDK4/6 inhibitor treatment eventually develop resistance, while several studies suggest that a proportion of patients never benefit from this combination at all^38, 39^. Most successful targeted therapies have complementary diagnostic biomarkers to identify responsive patient populations^40^. There are no such clinically available biomarkers for predicting positive response to CDK4/6 inhibitors. In this sense, CDK4/6 inhibitors have been identified as a targeted yet “biomarker-untargeted” therapy^12, 41^. Because CDK4/6 inhibitors are expensive, require chronic administration, and have many associated side effects, identifying predictive biomarkers for their use without unnecessary physical and financial burden to patients is a growing area of research^42–44^.

Ataxia-telangiectasia mutated (ATM)/Checkpoint kinase (CHK)2 is an upstream cell cycle checkpoint complex that controls CDK4/6 activity^45^. Failure of ER+ breast cancer cells to activate ATM/CHK2 in response to endocrine therapy through somatic loss of DNA repair proteins like MLH1 preserves CDK4/6 activity, inducing resistance to standard endocrine therapies^7, 41^. The ATM-dependent G1 checkpoint is the only cell cycle phase that is sensitive to extrinsic mitogenic cues such as hormones like estrogen^46^. It is not surprising, therefore, that germline mutation of *ATM* and/or *CHEK2* predisposes incidence of ER+ breast cancer^47, 48^ and mutations in *CHEK2* specifically associate with poor prognostic ER+ breast cancer diagnoses^47, 49^. As germline mutations in *ATM* and *CHEK2* are rare^47, 50, 51^, somatic loss of proteins like MLH1 that mediate activation of this checkpoint, which are far more common events in patient tumors^10, 35, 52, 53^, become critical drivers of endocrine treatment response. Yet, little is known about the mechanism by which MLH1 activates the ATM-CHK2 axis, and the role of missense mutations in *MLH1* in creating separation of function between mismatch repair and G1 checkpoint regulation activities. Below, using multiple model systems and patient data we show that sub-clonal cellular niches with dysregulated MLH1, detectable in 12-15% of ER+ breast cancer patients, seed resistance to endocrine therapy and responsiveness to CDK4/6 inhibitors. These findings suggest a potential predictive diagnostic for first-line combinatorial administration of endocrine therapy and CDK4/6 inhibitors in an at-risk subset of ER+ breast cancer patients.

## Materials and Methods

### Cell Lines, Mice, si/shRNA Transfection and Growth Assays

Cell lines were obtained from ATCC (2015) and maintained and validated as previously reported. Mycoplasma tests were performed on parent cell lines and stable cell lines December 2023 with Lonza Mycoalert Plus Kit (Cat# LT07-710) as per the manufacturer’s instructions. Transient transfection with mutant MLH1 constructs (purchased from TWIST Biosciences) and siRNA against CDK4/6 (purchased from Sigma-Aldrich) was conducted using JetPrime PolyPlus transfection reagent. Stable cell lines were maintained in the presence of specified antibiotics at recommended concentrations. Knockdown and expression were validated using western blotting. Growth assays were conducted using cytoSmart cell counter. Growth assay results were plotted as fold change normalized as specified. Three-dimensional growth assays were conducted over 6-8 weeks with twice weekly drug treatments using standard protocols. Images were captured when colonies were established and then treatment was administered, with images taken again every week. Fold change in area of colonies was calculated over time and represented as percent growth. Tumor growth assays *in vivo* were carried out by injecting 5-7 x10^6^ MCF7 cells into the L4 mammary fat pad/mouse. Mice for MCF7 experiments were 4-to-6- week athymic nu/nu female mice (from Sanford Burnham Prebys animal facility). Tumors were randomized for treatment at ∼100-150 mm^3^ volume for MCF7 xenografts. Tumors were harvested at <2 cm diameter and were embedded in paraffin blocks, OCT and snap frozen. All mouse experiments were performed in compliance with all relevant ethical regulations for animal testing and research, and all experiments conducted in the study received approval from Sanford Burnham Prebys (IACUC # 21-047).

### Inhibitors and Agonists

All drugs were maintained as stock solutions in DMSO, and stock solutions were stored at –80 and working stocks at –20° unless otherwise mentioned. 4-hydroxy tamoxifen (cat# H7904) and fulvestrant (cat# I4409) were purchased from Sigma-Aldrich, and stocks were diluted to 10 mmol/L working stocks for all experiments other than dose curves, where specified concentrations were used. For all experiments, cells were treated 24 hours after plating, and thereafter every 48 hours until completion of experiment. For mouse xenograft experiments, fulvestrant concentrations of 250 mg/kg body weight were used. Beta-estradiol was purchased from Sigma-Aldrich (cat# E8875), maintained in sterile, nuclease-free water, and diluted to obtain 10 mmol/L stocks for all experiments. For mouse xenograft experiments, 17 β-estradiol was added to the drinking water twice a week at a final concentration of 8 μg/mL (cat# E2758; Sigma). CHK2 activator (3, 3;-diindolyl methane, cat# sc-204624; Santa Cruz Biotechnology) was used at 100 μmol/L concentrations. Abemaciclib (LY2835219, cat# S7158; SelleckChem) and palbociclib (PD-0332991, cat# S1116; SelleckChem) were used at 2.5 μmol/L final concentrations for cell line assays and at 70 mg/kg/day in chow for tumor growth assays as described previously.

### Flow cytometry, Immunostaining and Microscopy

Flow cytometry was conducted to determine differential growth effects caused by presence or absence of estrogen (E_2_) for cells harboring WT or mutant *MLH1* when grown in co-culture. MCF7 cells with G55V mutation that were positive for GFP were plated at a 1:9 ratio with WT MLH1 MCF7 cells. Cells were analyzed at D0 and D7 post E_2_ or no E_2_ treatment. On D0 and D7 cells were trypsinized, washed and resuspended in 1x PBS + 2% FBS supplementation with DAPI as a viability marker and stored on ice until analyzed. GFP+ and GFP-populations of live cells were determined by flow cytometry using Aurora 5-laser Full Spectrum Analyzers (Aurora 5L). Immunostaining was performed based on manufacturer’s instructions. Cells were washed in 1x PBS; fixed for 20 minutes at room temperature in 4% PFA; blocked for 1 hour at room temperature in 5% goat serum and 1% Triton X-100 in 1x PBS; incubated with primary antibody overnight at 4 degrees in 1% goat serum and 1% Triton X-100 in 1x PBS antibody diluent; incubated with secondary antibody in diluent for 1 hour at RT; and then mounted with DAPI-containing mounting media (cat# P36935). Primary antibodies used include MLH1 (Cell Signaling Technology; cat# 4256; 1:200), pChk2 (cat# 2197), PCNA (cat# 2586) and Ki67 (cat# 9027). Fluorescent images were captured with an ECHO microscope and quantified with ImageJ.

### qPCR and Western Blotting

2×10^5^ cells were seeded overnight per well in 6-well plates. Following treatment with fulvestrant 1µM for 36 h, total RNA was extracted using RNeasy Mini Kit (Qiagen, Hilden, Germany). cDNA was synthesized using oligo(dT) and random primers (AB Bioscience, MA, USA), and qPCR analysis was performed with SYBR Green (Roche, NJ, USA). Primers for each were designed using NCBI primer blast and were ordered from Integrated DNA Technologies (IDT). 36B4 was used as internal control. Western blotting was conducted as previously described. All antibodies were purchased from Cell Signaling Technology and used at 1:1000 dilutions unless otherwise specified. Primary antibodies were incubated with the membrane overnight at 4 degrees and included MLH1 (cat# M8320), CDK6 (cat# D48S8) and GAPDH (cat# 2175, 1:5,000).

### 3D Spheroids and Sub-clonal Assays

For 3D spheroid assays, 4×10^3^ T47D cells were plated in 96 well-plate pre-coated with 1.5% agar. For sub-clonal 3D assay, G55V (GFP) and WT T47D cells were mixed in 1:9 ratio. Spheroids were allowed to grow for 1 week followed by respective treatments for 4 to 6 weeks. For -E_2_/+E_2_ experiments, 10nM of estradiol was added in the +E_2_ condition biweekly. For all other drug assays, fresh drug or vehicle were added biweekly. Spheroid pictures were captured using ECHO microscope and quantification of size was done using image J software. For *in vivo* sub-clonal xenograft experiment, G55V (GFP) and WT MCF7 cells were mixed in 1:9 ratio. A total of 5×10^6^ cells were injected into the left flank. Tumors were randomized for +E_2_ and -E_2_ group at ∼100-150 mm^3^ volume. Tumor measurements were taken biweekly and bioluminescence pictures of mice were also captured using IVIS spectrum imaging system. Tumors were harvested at <2 cm diameter and were embedded in paraffin blocks, OCT and snap frozen.

### PDxO Establishment and Treatments

For PDxO preparation, PDX tissue chunks were digested in a GentleMACS dissociator in warm advanced DMEM/F12 with GentleMACS human tumor dissociation enzymes and 10 μM Y-27632 added. After digestion, differential centrifugation was performed to enrich for organoids and deplete single cells. Organoids were embedded in 200-μl Matrigel domes, which were plated in six-well tissue culture plates onto a 50-μl Matrigel base layer. After a 5-min incubation period, plates were flipped, and Matrigel domes were solidified for 10 min before subtype-specific culture medium was added. For all breast cancer subtypes, 10 μM Y-27632 was added fresh to the PDxO base medium (Advanced DMEM/F12 with 5% FBS, 10 mM HEPES, 1× Glutamax, 1 μg ml–1 hydrocortisone, 50 μg ml–1 gentamicin and 10 ng ml–1 hEGF). Medium was exchanged every 3-4 days, and, once mature, cultures were passaged by incubating in dispase solution (20% FBS in dispase with Y-27632), followed by a wash step with base medium and a dissociation step in TrypLE Express. Single cells were seeded at 200,000– 400,000 cells per dome. For drug treatments, Mature organoids were collected from culture using dispase treatment at 37 °C for 25 min. Five thousand to ten thousand cells (∼50 organoids) were seeded per well in 96 tissue culture plates, each comprising a solidified 10 μl Matrigel base layer and 100 μl of PDxO medium supplemented with 5% (by volume) Matrigel. Media was replaced every 48 h with fresh drug. Cell titre glo 3D was conducted to measure viability.

### Categorization of Cytoplasmic vs Nuclear MLH1 in PDXs and Patient Tumors

PDXs were categorized as nuclear MLH1 if they had >5% cells with detectable nuclear localization of MLH1 and as cytoplasmic MLH1 if they had <5% cells with detectable nuclear localization of MLH1 but >10% of cells with detectable cytoplasmic localization of MLH1 (representative images in Fig S6). Patient tumor biopsies were pathologically graded for cytoplasmic MLH1 on a scale of 0 to 3; where a score of 0 represents no detectable signal for MLH1 in the cytoplasm and a score of 3 reflects >90% of cells with cytoplasmic MLH1 (representative images in Fig 5F). A similar scale was used to quantify nuclear presence of MLH1 in the same tumors. Tumors were deemed cytoplasmic MLH1 if they had detectable MLH1 in the cytoplasm (score of 1-3), and nuclear MLH1 if they had detectable MLH1 staining but no cytoplasmic MLH1 (score of 0). Tumors with no detectable MLH1 were deemed deficient for MLH1.

### Statistical Analysis

Statistical significance between two groups was assessed by the unpaired Student’s t-test. For spheroid experiments and tumor growth rates, slopes were calculated and compared for statistical differences. For categorical data, a Fisher’s Exact or a Pearson’s Chi-squared test was used as indicated based on sample size. All experiments were conducted in triplicate and each experiment was duplicated ≥2 times.

## Results

### *MLH1* Mutations Drive Varying Degrees of Endocrine Treatment Resistance in ER+/HER2 Breast Cancer Cells

We identified 6 somatic *MLH1* mutations from primary and metastatic ER+/HER2- breast cancer patient samples in three independent datasets (NeoPalAna (primary), TCGA (primary) and MSKCC (primary and metastatic)). Three of the selected mutations were missense (G55V, E199K and D584H) and three were nonsense (E414*, E439* and E717*) (Fig 1A). We then transfected two ER+/HER2- breast cancer cell lines (MCF7 and T47D) stably expressing shRNA against *MLH1* (shMLH1) with sh-resistant constructs of each mutant *MLH1* cDNA or the wildtype. Protein levels of missense mutated *MLH1* constructs are comparable to that of wildtype constructs in both transiently (Fig 1B) and stably transfected (Fig S1A) cells.

**Figure 1:**
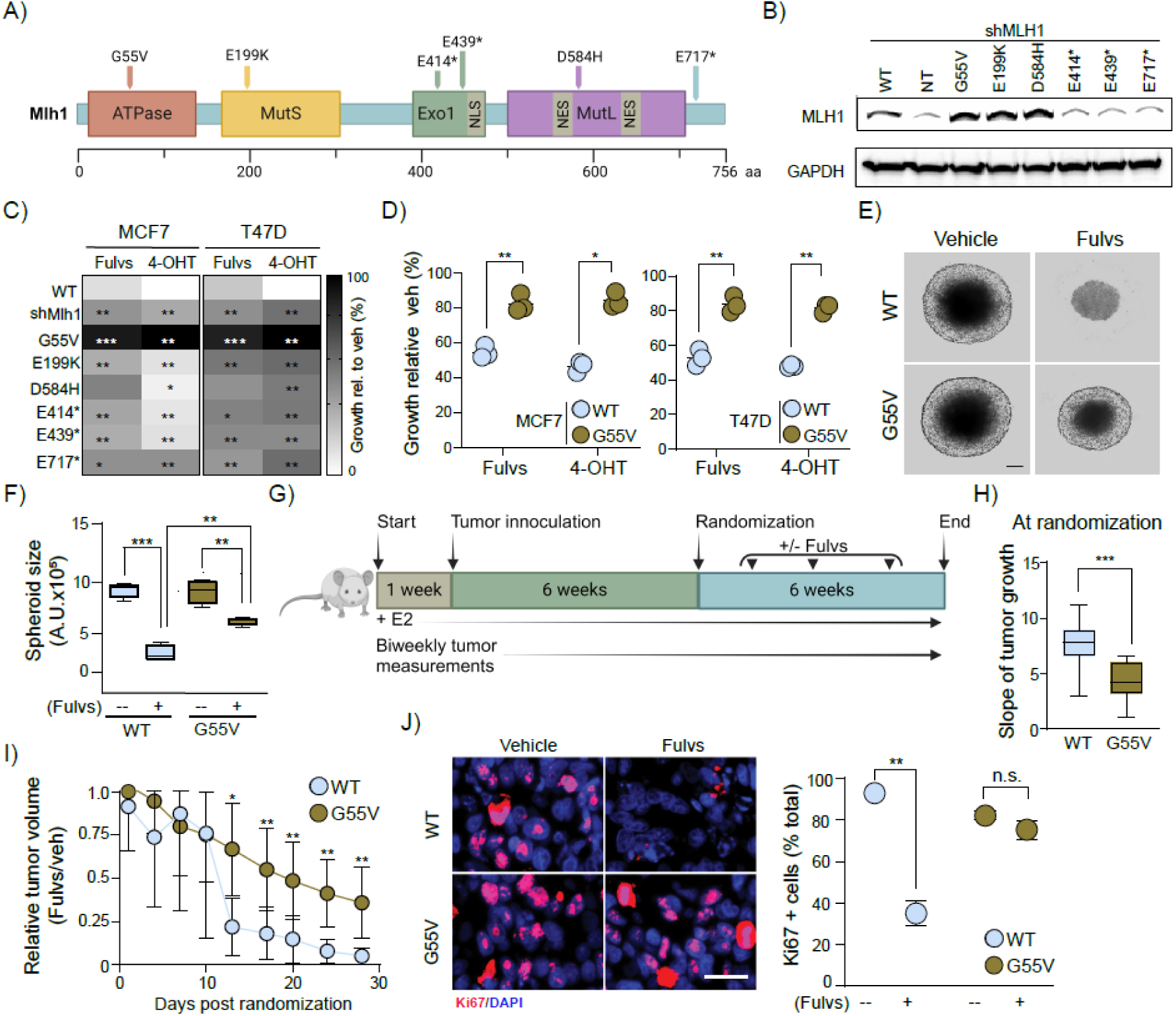
Some *MLH1* mutations induce endocrine therapy resistance. (A) Schematic showing mutations in *MLH1* from patient tumors. (B-C) Western blot assessing levels of MLH1 protein (B) and heatmap indicating growth of cells treated with specific endocrine therapy (100nM fulvestrant or 4-OHT) (C) after transient expression of sh-resistant mutant and wild type constructs in MCF7 cells stably expressing shRNA against *MLH1*. Associated data validating endocrine therapy response in MCF7 and T47D cells stably expressing mutants presented in Fig S1A-B. (D) Dot plots depicting growth of MCF7 and T47D cells of indicated genotypes after endocrine treatment relative to vehicle. (E-F) Representative images of T47D^G55V^ and T47D^WT^ spheroids after fulvestrant treatment (E) and accompanying quantification (F). Associated data of 3D growth assays after 4-OHT treatment presented in Fig S1E. Lines in boxplots represent median. (G) Schematic of *in vivo* experimental design. (H-I) Boxplot depicting differences in tumor growth rate at baseline before randomization (calculated as slope of volume over time for each tumor) (H) and tumor volume normalized to vehicle controls with biweekly administration of fulvestrant (I) to mice transplanted with MCF7^G55V^ or MCF7^WT^ cells. Raw tumor volumes presented in Fig S1F. (J) Representative images of Ki67 staining (proliferation marker, red) in MCF7^G55V^ or MCF7^WT^ tumors (DAPI as nuclear stain) with and without fulvestrant treatment with accompanying quantification. Student’s t-test determined p-values. *** represents p≤0.0001, ** p≤0.01 and * p≤0.05. For all graphs, error bars describe standard deviation. Scale bar = 20µm. Abbreviations: Vehicle (Veh), Fulvestrant (Fulvs), Tamoxifen (4-OHT), Wild type (WT), not significant (n.s.)

On the other hand, as expected, cell lines with nonsense mutated *MLH1* constructs express lower levels of MLH1 analogous to shMLH1 cells likely because they undergo nonsense mediated decay (Fig 1B, Fig S1A). Next, we tested whether these selected *MLH1* mutations alter response to endocrine therapy. We treated MCF7 and T47D cells with transient (Fig 1C), and T47D cells with stable (Fig S1B), expression of the six mutant constructs with two common classes of endocrine therapy: fulvestrant or tamoxifen and assessed 2D growth. We observed no significant difference in baseline growth for any of these mutations whether expressed in MCF7 (Fig S1C) or T47D (Fig S1D) cells. However, in shMLH1 cells transfected either transiently (Fig 1C) or stably (Fig S1B) with each of the 6 mutant *MLH1* constructs, we observed varying levels of endocrine treatment resistant relative to isogenic cells with MLH1^WT^. As expected, control isogenic cells with stable expression of shMLH1 demonstrate robust endocrine treatment resistant growth relative to shMLH1 cells transfected with wildtype *MLH1* (MLH1^WT^) constructs (Fig 1C: transient, Fig S1B: stable). Of all the mutations, G55V, a missense mutation in the ATPase domain of *MLH1*, most significantly and consistently induces resistance to endocrine treatment, even surpassing shMLH1 cells (Fig 1C: transient, Fig S1B: stable).

Both MCF7^G55V^ and T47D^G55V^ cells continue to grow at 85-90% efficiency of vehicle-treated controls in presence of either type of endocrine treatment unlike MCF7^WT^ and T47D^WT^ counterparts which fall to ∼50% growth efficiency after endocrine therapy (Fig 1D). Since *MLH1^G55V^* drives endocrine treatment resistance in two cell lines, we studied its effect on 3D growth using T47D^G55V^ cells (stable expression).

T47D^G55V^ spheroids retain ∼60% 3D growth efficiency after fulvestrant treatment compared to T47D^WT^ spheroids which fall to ∼35% growth (Fig1E-F; p=0.0004). These results are reproducible with tamoxifen treatment (Fig S1E; p=0.00005). To validate these findings *in vivo*, we xenografted MCF7^G55V^ or MCF7^WT^ cells (stable expression) into mammary fat pads of nude mice (Fig 1G). Of note, in untreated mice, MCF7^G55V^ tumors grow slowly, at 57% of the rate of MCF7^WT^ tumors (Fig 1H; p=0.0001). However, MCF7^G55V^ tumors appear more resistant to fulvestrant treatment, remaining at 35% the size of vehicle-treated counterparts compared to MCF7^WT^ tumors which show complete regression to ∼3% (Fig 1I; p=0.007; raw tumor volume data in Fig S1F). In keeping with fulvestrant resistant tumor growth *in vivo*, MCF7^G55V^ tumors from fulvestrant-treated mice continue proliferating at the same rate as those from vehicle-treated mice, unlike MCF7^WT^ tumors, which retain only 27% of their proliferative capacity (p=0.001), as assessed by Ki67 positivity (Fig 1J).

Next, to test whether endocrine treatment resistance induced by *MLH1^G55V^* is due to loss of dependence on estrogen signaling for growth, we assayed growth in response to estrogen deprivation in serum-starved media. In MCF7^WT^ cells, estrogen deprivation decreases growth to 22% of that observed in estrogen-supplemented cells, while MCF7^G55V^ cells retain 61% growth efficiency (Fig 2A; p=0.003). In T47D^WT^ cells, similarly, estrogen deprivation decreases growth to 4% of that seen in estrogen-supplemented counterparts, while T47D^G55V^ cells remain at 19% growth efficiency (Fig 2B; p=0.003). As expected both MCF7^G55V^ and T47D^G55V^ cells remain resistant to both fulvestrant and tamoxifen even in serum starved media where estrogen is the primary growth stimulus unlike MCF7^WT^ and T47D^WT^ counterparts (Fig 2A-B).This decreased dependence on estrogen receptor signaling is even more pronounced in 3D growth assays where estrogen deprivation has little to no detectable impact on T47D^G55V^ (p=0.3) spheroids while T47D^WT^ spheroids decrease dramatically (p=0.00003) to ∼22% of the size of estrogen-supplemented controls (Fig 2C). These observations validate *in vivo*, where estrogen deprivation of mice xenografted with MCF7^G55V^ cells has no effect on tumor growth relative to estrogen-supplemented controls (Fig 2D-E, p=0.3). Together, these findings support a causal role for MLH1^G55V^ in endocrine treatment resistance and estrogen-independent growth despite robust and stable expression of the mutant protein (shown in Western blots in Fig S1A).

**Figure 2:**
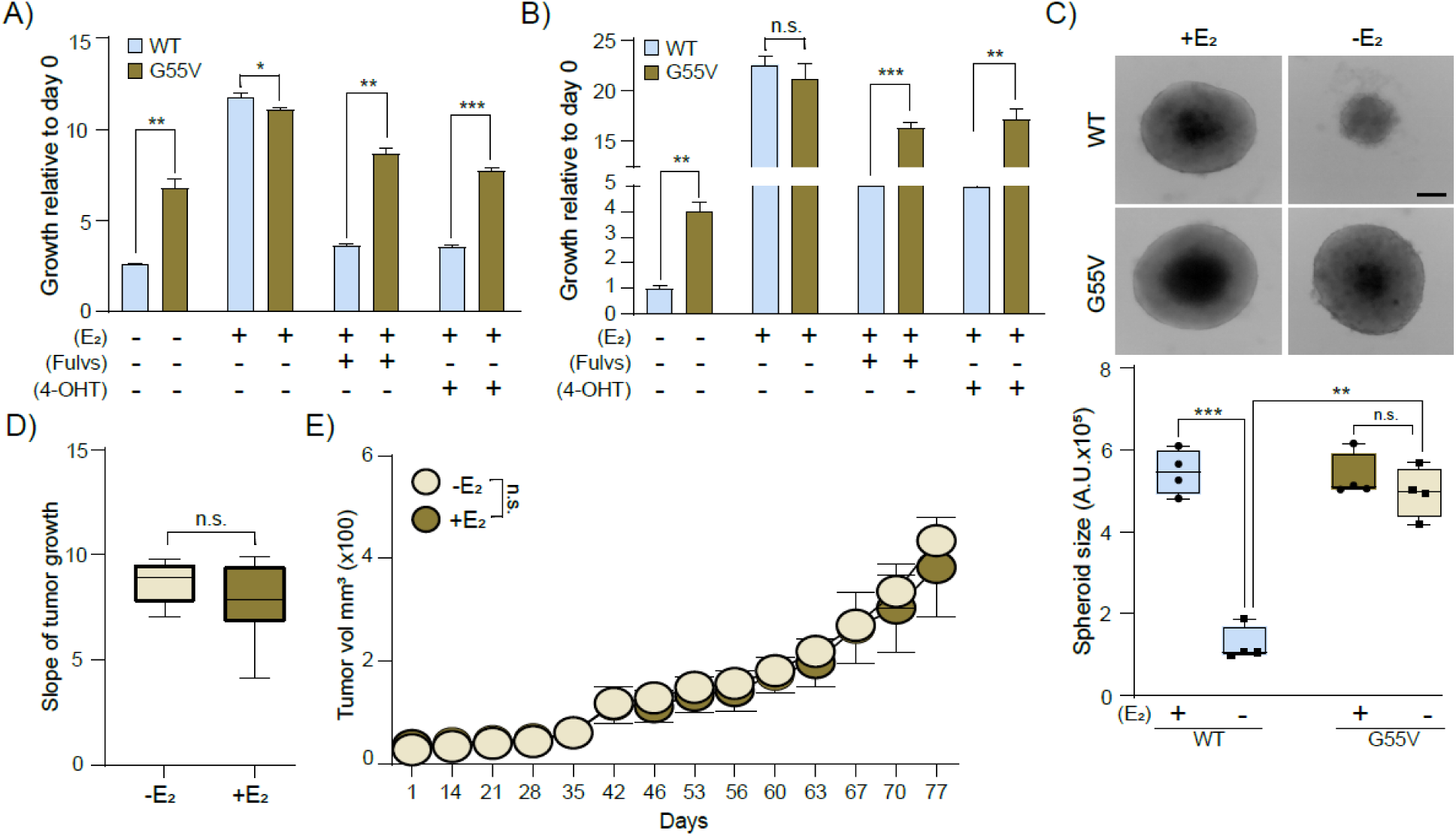
Some *MLH1* mutations induce estrogen-independent growth of ER+ breast cancer cells. (A-B) Bar graph depicting growth of MCF7 (A) and T47D (B) cells of specified genotypes at day 7 relative to day 0 with or without addition of 10nM beta-estradiol to cells grown in charcoal stripped serum and treated with Fulvs (100nM) or 4-OHT (100nM) as indicated. (C) Representative images of 3D spheroids of T47D cells grown in charcoal stripped serum with (+E_2_) or without (-E_2_) 10nM beta-estradiol treatment with accompanying box plots showing quantification of spheroid size. (D-E) Boxplots depicting differences in tumor growth rate (calculated as slope of volume over time for each tumor) of MCF7^G55V^ cells with or without estrogen supplementation in drinking water (D) with accompanying line graphs of average tumor volume over time for indicated groups (E). Student’s t-test determined p values. Scale bar = 50µm. *** represents p≤0.0001, ** p≤0.01 and * p≤0.05. Abbreviations: estrogen (E_2_), Fulvestrant (Fulvs), Tamoxifen (4-OHT), Wild type (WT), Not significant (n.s.).

### Sub-clonal Cell Populations with *MLH1* Mutation Seed Endocrine Treatment Resistance

We originally identified *MLH1^G55V^* in a primary ER+/HER2- breast cancer patient sample at a low VAF of 3.38% (Fig 3A, inset). After treatment with anastrozole (an aromatase inhibitor, with a similar mechanism of action to estrogen deprivation) for 4 weeks this patient was highly responsive to treatment with her tumor demonstrating 62% decrease in proliferating cells, assessed using immunohistochemistry for proliferation marker Ki67 (Fig 3A). This proliferative inhibition in response to endocrine therapy is comparable to that of other endocrine therapy responsive patients in the trial whose tumors had no detectable mutation in/dysregulation of *MLH1* (Fig 3A). These findings appear to contradict our experimental data *in vitro* and *in vivo* showing a causal role for *MLH1^G55V^* in inducing estrogen-independent growth (Fig 2) and endocrine treatment resistance (Fig 1). However, initial reduction in bulk tumor size may not be predictive of ultimate outcome, since it cannot detect treatment response of sub-clonal *MLH1^G55V^* cells within the tumor that may seed recurrence.

**Figure 3:**
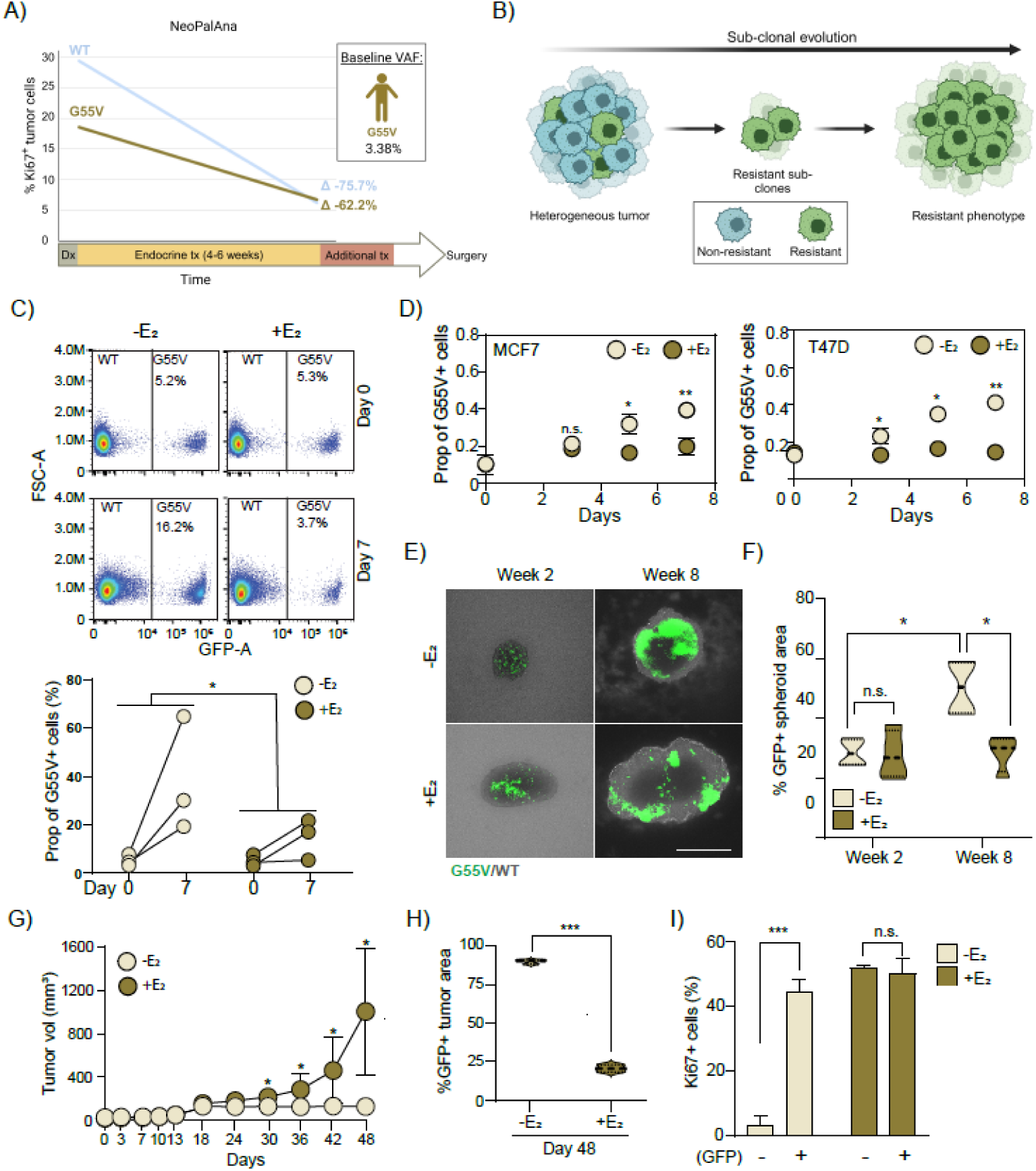
*MLH1^G55V^* cells persist in sub-clonal fractions after administration of endocrine therapy. (A) Schematic from NeoPalAna trial showing variant allele fraction of MLH1^G55V^ (inset) in a patient tumor and tumor proliferative response to endocrine therapy assessed using Ki67. (B) Schematic of sub-clonal evolution. (C) Representative images from flow cytometry assay showing MCF7^G55V^ (GFP+) and MCF7^WT^ (untagged) cells mixed in 1:9 ratio and grown with or without 10nM of beta-estradiol for a week with accompanying quantification. (D) Dot plots quantifying proportions of GFP+ MCF7^G55V^ and T47D^G55V^ lines grown with indicated treatments. Associated data showing response to fulvestrant and tamoxifen presented in Fig S2A-D. (E-F) Representative images of 3D spheroids (E) of T47D^G55V^ and T47D^WT^ cells admixed at a 1:9 ratio grown with or without 10nM of beta-estradiol for 8 weeks with accompanying quantification presented as a violin plot (F). Associated data showing overall spheroid size and response to fulvestrant and tamoxifen are presented in Fig S2E-G. (G) Average tumor volume of MCF7^G55V^ (GFP+) and MCF7^WT^ cells mixed in 1:9 ratio transplanted into mice with or without estrogen supplementation in drinking water. (H) Proportion of MCF7^G55V^ (GFP+) cells in each tumor at harvest based on quantification of GFP+ cells using immunofluorescence. (I) Quantification of Ki67+/GFP+ (*MLH1^G55V^*) and Ki67+/GFP- (*MLH1^WT^*) cells in indicated conditions. Representative images presented in Fig S3. Student’s t- test determined all p values. *** represents p≤0.0001, ** p≤0.01 and * p≤0.05. For all graphs, error bars describe standard deviation. Scale bar = 100µm. Abbreviations: estrogen (E2), Wild type (WT), not significant (n.s.).

To test this hypothesis, we experimentally replicated the sub-clonality observed in patient tumors by admixing untagged MCF7^WT^/T47D^WT^ and GFP-tagged MCF7^G55V^/T47D^G55V^ isogenic cells at a 9:1 ratio, respectively (Fig 3B). In flow cytometry assays, MCF7^G55V^ cells are ∼4-fold enriched for viability over MCF7^WT^ cells after estrogen deprivation but show no enrichment in estrogen-supplemented control conditions (Fig 3C; p=0.02). We confirmed these findings with orthogonal 2D growth assays in both MCF7 and T47D cells where we immunohistochemically quantified GFP+ cell populations over 7 days of growth in response to estrogen deprivation relative to estrogen-supplemented control arms. Similar to the flow cytometry results, MCF7^G55V^ and T47D^G55V^ cells grow from 10% to constitute 50% of the total cell population in estrogen-deprived conditions but remain at 10% sub-clonality over time in estrogen-supplemented control groups (Fig 3D; MCF7: p=0.004; T47D: p=0.01). These data suggests that G55V-bearing cells have little or no growth advantage intrinsically but assume adaptive growth advantage when exposed to endocrine interventions. We confirmed these observations in response to fulvestrant (MCF7: Fig S2A, T47D: Fig S2C) and tamoxifen (MCF7: Fig S2B, T47D: Fig S2D) treatment. These findings also validate in 3D assays. After 6 weeks of estrogen deprivation, overall spheroid size significantly decreases relative to estrogen-supplemented controls (Fig S2E), but the proportion of T47D^G55V^ cells more than doubles (p=0.01) in estrogen-deprived conditions while remaining constant in estrogen-supplemented controls (Fig 3E-F). Similarly, the T47D^G55V^ proportion increases 2-fold with fulvestrant (p=0.00008) and 3-fold with tamoxifen (p=0.000003) treatment, relative to vehicle controls (Fig S2E-F). We then replicated sub-clonality *in vivo* by xenografting mixed populations of MCF7^WT^ (untagged): MCF7^G55V^ (GFP-tagged) cells at a 9:1 ratio to form tumors in the mammary fat pads of nude mice. Mice were initially supplemented with physiological levels of estradiol until tumors were large enough to randomize into treatment groups: estrogen-deprived or continued estrogen supplementation. As expected, estrogen deprivation significantly decreases bulk tumor volume (p=0.01) relative to estrogen-supplemented controls (Fig 3G). However, the proportion of MCF7^G55V^ cells (assessed by quantification of the GFP tag) increases to ∼100% of the tumor composition in estrogen-deprived mice, while remaining at ∼20% tumor composition in the estrogen-supplemented control arm (Fig 3H, p=0.00001).

These observations are supported by assessment of proliferation in the tumors, where MCF7^G55V^ tumor cells in estrogen-deprived mice maintain proliferation at rates comparable to MCF7^G55V^ tumor cells in estrogen-supplemented control mice (p=0.52), while MCF7^WT^ counterparts within the same tumor significantly inhibit proliferation (p=0.0002) in estrogen-deprived vs -supplemented mice (Fig 3I, representative images: Fig S3). These findings together suggest that *MLH1^G55V^* cells in patient tumors may escape standard first-line endocrine monotherapy. Because they are sub-clonal, however, they likely remain undetected for extended periods of time while seeding recurrence.

### G55V Mutation Reduces Chk2 Activation Leading to Endocrine Therapy Resistance

To identify the mechanism by which *MLH1^G55V^* confers endocrine therapy resistance, we conducted RNAseq analysis of isogenic MCF7^WT^ and MCF7^G55V^ cells at baseline and after fulvestrant treatment. We included vehicle- and fulvestrant-treated MCF7 shMLH1 and MCF7^E717*^ cells as additional controls to identify pathways uniquely dysregulated by *MLH1^G55V^*. Pathway analysis identified cell cycle dysregulation (Fig S4A), specifically G1/S checkpoint activity (Fig 4A), as a distinguishing feature of the MCF7^G55V^ cellular response to fulvestrant treatment (FDR<0.05). Chk2, a key G1 checkpoint kinase, is one of the top cell cycle regulators downregulated in MCF7^G55V^ relative to control cells (Fig 4A). To validate this RNAseq observation, we assessed cells for nuclear, phosphorylated (therefore, active) Chk2 protein expression in our experimental model systems. While MCF7^WT^ cells register a 16-fold increase in cells with detectable nuclear pChk2 positivity upon fulvestrant treatment (p=0.004), MCF7^G55V^ cells have no detectable increase (Fig 4B). These results validate in T47D cells as well (Fig S4B). Since Chk2 activation is known to arrest G1-S transition when ER+/HER2- breast cancer cells are exposed to endocrine therapy resulting in a proliferative block^7, 54^, we also assessed expression of S phase marker PCNA in MCF7^G55V^ and MCF7^WT^ xenograft tumors grown in mice treated with either vehicle or fulvestrant. MCF7^WT^ tumors have ∼70% decrease in S-phase (PCNA+) cells in response to fulvestrant treatment, while MCF7^G55V^ tumors have a ∼20% decrease (Fig 4D; p=0.006). This striking discrepancy in Chk2 activation and associated G1-S transition suggests that impaired Chk2 activation in *MLH1^G55V^* tumors leading to a failed G1-S checkpoint block in response to endocrine treatment drives treatment resistance.

**Figure 4:**
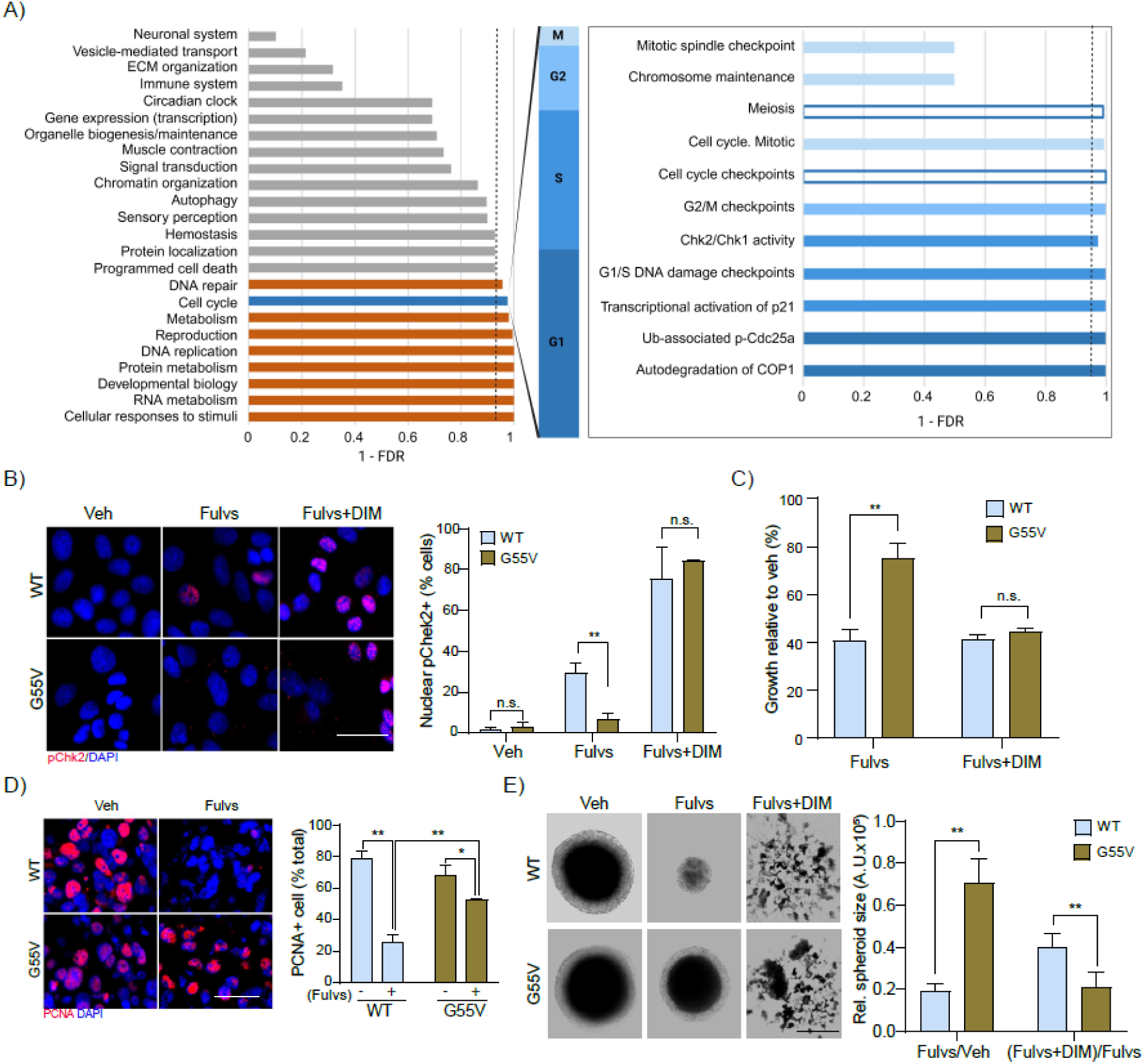
*MLH1^G55V^* prevents Chk2 activation despite abundant and stable expression. (A) Gene set enrichment analysis of RNAseq data comparing MCF7^G55V^ to MCF7 shMLH1, MCF7^WT^ and MCF7^E717^* cells after treatment with 1 µM fulvestrant for 36h with bar graphs depicting the significance of individual cell cycle component pathways (expressed as 1-FDR for visibility). Associated Reactome analyses are presented in Fig S4A. (B) Representative immunofluorescence photomicrographs showing pChk2 sub-cellular localization in MCF7^WT^ and MCF7^G55V^ cells with or without 36h of treatment with 1µM fulvestrant alone or a combination of fulvestrant (100nM) + DIM (200µM). Accompanying quantification is presented as bar graphs. Associated data showing pChk2 expression in T47D cells are presented in Fig S4B-C. (C) Bar graphs showing growth of MCF7^WT^ and MCF7^G55V^ cells after indicated treatments for 36h. Growth is normalized to vehicle for each type and expressed as percent inhibition. Associated data showing response in T47D cells is presented in Fig S4D. (D) Representative immunofluorescence photomicrographs showing PCNA expression in MCF7^WT^ and MCF7^G55V^ xenograft tumors with indicated treatment with accompanying quantification. (E) Representative images of T47D^WT^ and T47D^G55V^ spheroids after biweekly treatment with 10nM fulvestrant alone or with a combination of fulvestrant+DIM (10nM+50µM) with accompanying quantification. Student’s t-test determined all p-values. *** represents p≤0.0001, ** p≤0.01 and * p≤0.05. For all graphs, error bars describe standard deviation. Scale bar = 20µm for (B and D) and 100µm for (E). Abbreviations: Vehicle (Veh), Fulvestrant (Fulvs), 3,3′-Diindolylmethane (DIM).

To test this hypothesis functionally, we used small molecule Chk2 activator, di-indolyl methane (DIM) to exogenously activate Chk2 in MLH1^G55V^ cells. DIM, unlike fulvestrant, treatment induces robust Chk2 activation comparable to that induced by fulvestrant in MLH1^WT^ cells (MCF7: Fig 4B, T47D: Fig S4C). In validation of our hypothesis that blunted Chk2 activation mediates endocrine treatment resistance in MLH1^G55V^ cells, administration of DIM rescues sensitivity to fulvestrant in 2D growth assays (MCF7: Fig 4C, T47D: Fig S4D). This observation also validates in 3D assays where addition of DIM to fulvestrant decreases spheroid size of T47D^G55V^ cells by 71% when compared to fulvestrant alone, phenocopying the effect of fulvestrant monotherapy on T47D^WT^ cells (Fig 4E; p=0.0003).

### Aberrant Subcellular Localization of MLH1 Dampens Chk2 Activation in Response to Treatment

We, and others, have shown that loss of MLH1 mutes Chk2 activation, an event critical for mounting cell cycle arrest in response to endocrine treatment^7, 41, 55, 56^. However, the MLH1^G55V^ protein is abundantly and stably expressed in ER+/HER2- breast cancer cell lines as detected by Western blotting (transient expression: Fig 1B, stable expression: Fig S1A), suggesting that loss of Chk2 activation in this case is not due to destabilization of the MLH1 protein. Therefore, we next assessed subcellular localization of MLH1^G55V^ using immunofluorescence. Upon endocrine treatment, MCF7^WT^ and T47D^WT^ cells show a striking increase in cells with nuclear localization of MLH1 compared to the vehicle control arm (Fig 5A: MCF7; p=0.0008; Fig S5: T47D; p=0.01), consistent with previous reports^7^. However, cells expressing MLH1^G55V^ show no detectable increase in MLH1 nuclear translocation, remaining cytoplasmic both before and after endocrine treatment (MCF7: Fig 5A, T47D: S5). Lack of nuclear MLH1 persists *in vivo* in xenografted MCF7^G55V^ tumors in contrast to MCF7^WT^ counterparts (Fig 5B; p=0.009). This loss of nuclear localization, therefore, likely phenocopies the effect of complete MLH1 loss in terms of Chk2 activation.

**Figure 5:**
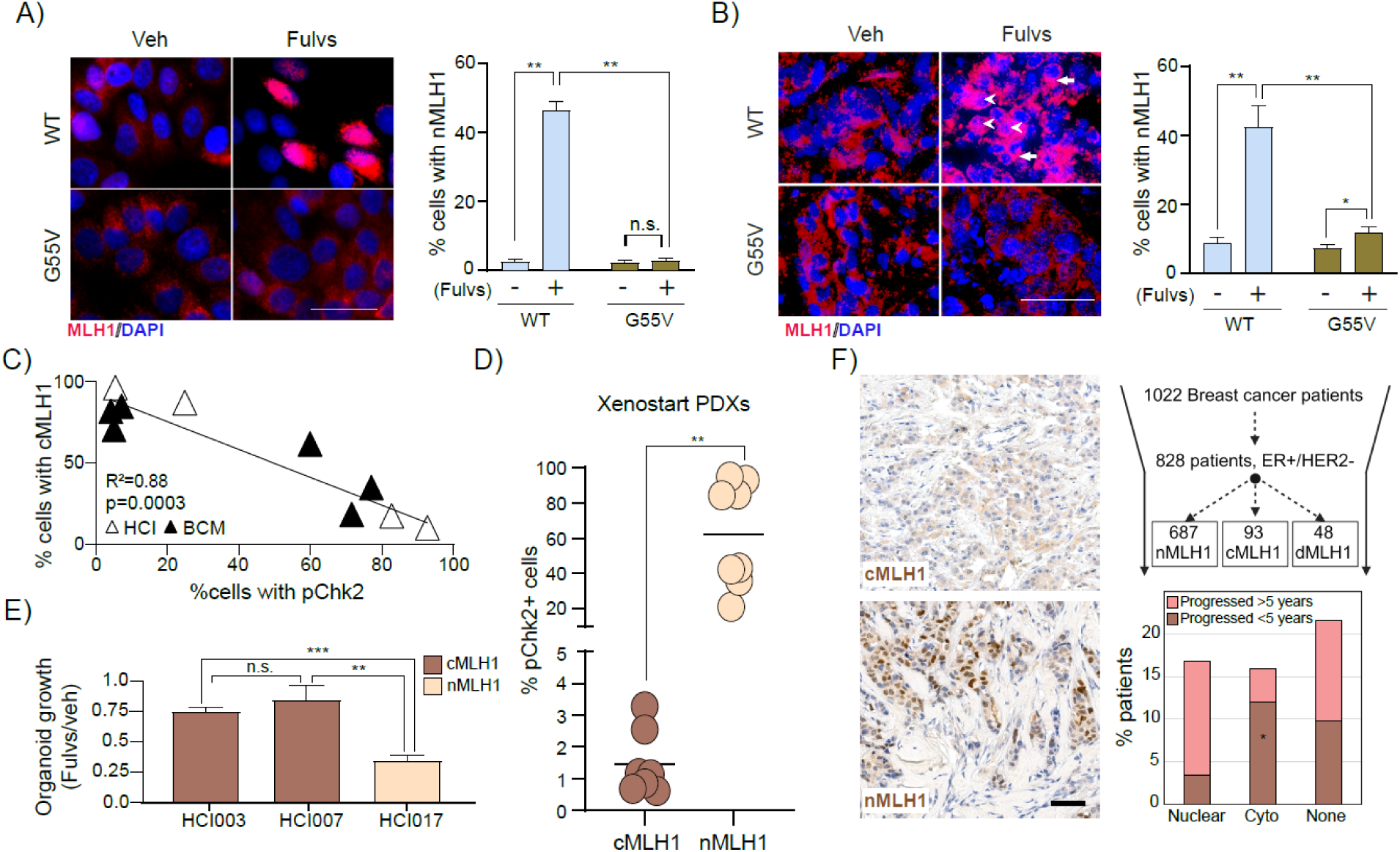
G55V alters subcellular localization of MLH1. (A-B) Representative immunofluorescence images showing subcellular expression of MLH1^WT^ vs MLH1^G55V^ in MCF7 cells (A) and xenograft tumors (B) after fulvestrant treatment. Accompanying bar graphs represent quantification of percent cells with detectable nuclear MLH1. Data for subcellular localization of MLH1^WT^ and MLH1^G55V^ in T47D cells is presented in Fig S5. Arrowheads in panel B indicate cells with nuclear MLH1 and arrows, cytoplasmic MLH1. (C) Regression analysis showing association between cytoplasmic expression of MLH1 and nuclear pChk2 positivity in HCI and BCM PDX lines. (D) Dot plot representing quantification of cells positive for cytoplasmic localization of MLH1 and nuclear localization of pChk2 in each ER+/HER2- PDX tumor of the Xenostart panel. Associated representative images of MLH1 and pChk2 expression for all PDXs is presented in Fig S6. (E) Bar graphs quantifying percent cell viability of three PDX derived organoids models after fulvestrant treatment relative to vehicle-treated controls. (F) Categorial analysis performed on patient data. Fisher’s Exact test determined p-value. Associated information on patient tumor dataset depicted in Fig S7. Student’s t-test determined all p values except for panel F. *** represents p≤0.0001, ** p≤0.01 and * p≤0.05. For all graphs, error bars describe standard deviation. Scale bars = 50µm (F). Abbreviations: Vehicle (Veh), Fulvestrant (Fulvs), Wild type (WT), Huntsman Cancer Institute PDXs (HCI), Baylor College of Medicine PDXs (BCM), not significant (n.s.), cMLH1 (cytoplasmic MLH1), nMLH1 (nuclear MLH1).

To test this hypothesis, we investigated whether cytoplasmic localization of MLH1, independent of G55V mutation, prevents Chk2 activation to drive endocrine treatment resistance. We screened panels of MLH1^WT^ ER+/HER2- breast cancer patient-derived xenografts (PDXs) from three collections (Huntsman Cancer Institute (HCI), Baylor College of Medicine (BCM) and Xenostart) for subcellular localization of MLH1 and pChk2 using immunofluorescence.

First, combining HCI and BCM panels (n=4 HCI; n=6 BCM), we conducted an agnostic regression analysis between percent cells with cytoplasmic (but not nuclear) MLH1 and percent cells with nuclear pChk2 in each PDX tumor. In this analysis, cytoplasmic MLH1 presence inversely correlates with pChk2 positivity across PDX lines (Fig 5C with representative images from HCI: Fig S6A and BCM: Fig S6B). We independently validated this correlation in a third ER+/HER2- PDX panel (Xenostart, n=16). We categorized PDXs from this panel as nuclear or cytoplasmic MLH1 as described in Materials and Methods (representative images in Fig S6C). We found that nuclear MLH1 PDXs have ∼60% nuclear pChk2 positivity, while cytoplasmic MLH1 PDXs show <2% positivity for nuclear pChk2 (Fig 5D; p=0.0007). To test whether this decrease in Chk2 activation in PDXs with cytoplasmic MLH1 impacts their ability to respond to endocrine treatment, we conducted *ex vivo* organoid growth assays. Both HCI003 and HCI007 organoids (cytoplasmic MLH1) are resistant to endocrine therapy showing <25% growth inhibition after administration of fulvestrant, whereas HCI017 organoids (with nuclear MLH1) show strong responsiveness with 60% growth inhibition after endocrine treatment (Fig 5E; p=0.00009).

To test whether these associations between cytoplasmic MLH1 and endocrine therapy resistance are detectable in patients, we analyzed MLH1 subcellular localization and disease progression in a cohort of 1022 breast cancer patients, 828 of whom had ER+/HER2- disease (Fig S7A). Biopsies were pathologically categorized as nuclear or cytoplasmic MLH1 as described in Materials and Methods (Fig 5F for representative images). In this dataset ∼11% of all tumors have cytoplasmic MLH1 when including all subtypes, and ∼6% are deficient for MLH1, with the remaining 83% having detectable nuclear MLH1 (Fig S7B). These overall frequencies are comparable to that seen for ER+/HER2- breast cancer patient tumors alone (∼12% with cytoplasmic MLH1 and 5% deficient for MLH1) (Fig S7B). However, ER+/HER2- patient tumors are four times as likely to have high levels of cytoplasmic MLH1 (score ≥ 2) as HER2+ and triple negative tumors combined (Fig S7C-E). Of interest, cytoplasmic MLH1 presence in ER+/HER2- breast cancer patient tumors correlates with decreased time to progression (p=0.03), with 12% of patients with cytoplasmic MLH1 progressing on endocrine treatment within 5 years of diagnosis compared to only 2% of patients with nuclear MLH1 (Fig 5F). This increased immediate progression on first-line therapies is similar to that observed in MLH1 deficient patients (Fig 5F). Together these data strongly support a role for cytoplasmic MLH1 in mediating cell cycle dysregulation that precludes response to standard endocrine therapies in ER+/HER2- breast cancer patients.

### Cytoplasmic MLH1 Induces CDK4/6 Dependence in ER+/HER2- Breast Cancer Cells

ER+/HER2- breast cancer cells with cytoplasmic MLH1 fail to activate G1 checkpoint kinase Chk2 in response to endocrine treatment unlike their counterparts with robust nuclear translocation of MLH1 (Fig 6A).

**Figure 6:**
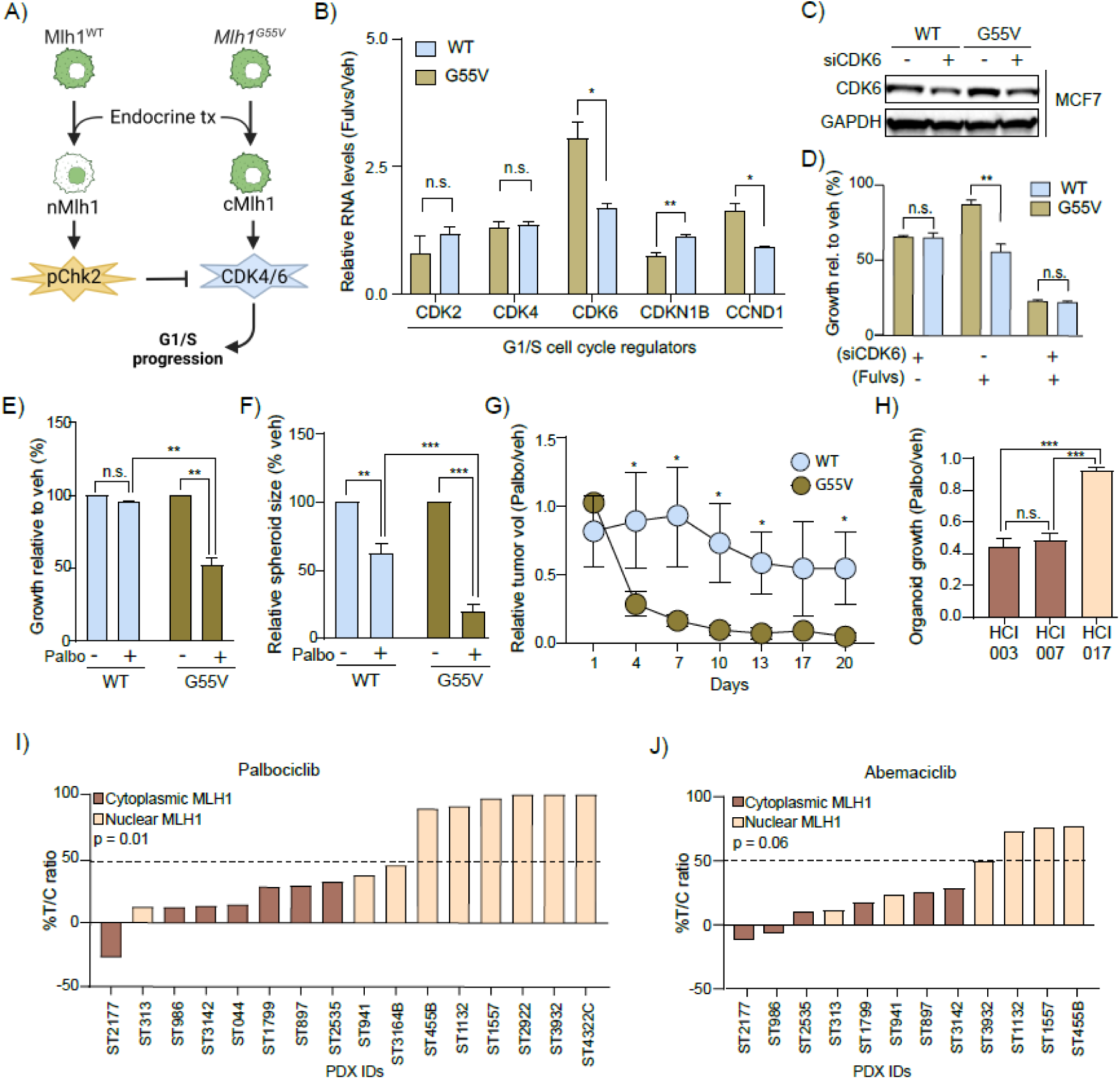
Cytoplasmic MLH1 predisposes sensitivity to CDK4/6 inhibition. (A) Schematic. (B) qRT-PCR analysis of MCF7 cells of indicated genotypes for relative expression of specified genes following fulvestrant treatment. (C-D) Bar graph showing growth of MCF7^WT^ and MCF7^G55V^ cells (D) after CDK6 knockdown (validated by Western blotting in C) alone or with fulvestrant treatment. Associated data representing growth in T47D cells in Fig S8A-B. (D) Bar graphs showing growth of MCF7^WT^ and MCF7^G55V^ cells after CDK4/6 inhibitor treatment. Associated data for T47D cells presented in Fig S8C and with abemaciclin in Fig S8D-E. (E) 3D growth of T47D^WT^ and T47D^G55V^ after indicated treatments. (F) *In vivo* tumor growth expressed as change in tumor volume of MCF7^WT^ and MCF7^G55V^ cells after palbociclib treatment normalized to vehicle. Associated data showing raw tumor volumes is presented in Fig S8F-G. (G) Bar graph showing effect of palbociclib treatment in three PDoX models. Growth is expressed relative to vehicle for each PDoX. (H-I) Waterfall plots depicting CDK4/6 inhibitor response expressed as percent growth of treated cells relative to vehicle-treated controls in PDX lines in response to (H) palbociclib (I) abemaciclib. Student’s t test determined all p values except for panels H-I where Fisher’s exact test was performed. *** p≤0.0001, ** p≤0.01 and * p≤0.05. For all graphs, error bars describe standard deviation. Abbreviations: Cyclin dependent kinases (CDK), Patient derived xenograft (PDX), Patient derived organoid (PDoX), Wild type (WT), Cyclin D1 (CCND1), Cyclin dependent kinase inhibitor 1B (CDKN1B).

Using MCF7^G55V^ cells to represent cells with aberrant cytoplasmic MLH1 localization, we next tested whether aberrant cytoplasmic localization of MLH1 alters other key G1/S-phase cell cycle regulators. Gene expression of G1 cyclin-dependent kinase *CDK6* increases 3-fold (p=0.01), and early G1 cyclin *CCND1* 1.6-fold in MCF7^G55V^ cells upon endocrine treatment (p=0.01) compared to MCF7^WT^ cells (Fig 6B). Additionally, MCF7^G55V^ cells downregulate *CDKN1B* the G1 inhibitor directly downstream of Chk2^57^, relative to MCF7^WT^ cells (Fig 6B; p=0.003). To test whether upregulation of G1 CDKs by cytoplasmic MLH1 induces cellular dependence on these CDKs for endocrine therapy resistant growth, we knocked down *CDK6* in MCF7^G55V^ (knockdown validated in Fig 6C) and T47D^G55V^ (knockdown validated in Fig S8A) cells using siRNA.

Knocking down *CDK6* in these cells restores sensitivity to endocrine treatment (MCF7: Fig 6D, T47D: Fig S8B), suggesting that aberrant cytoplasmic localization of MLH1 induces dependence on G1 CDKs for endocrine therapy resistance in ER+/HER2- breast cancer cells. We next tested whether CDK6 dependence in *MLH1^G55V^* cells results in sensitivity to CDK4/6 inhibitors. Both MCF7^G55V^ and T47D^G55V^ cells show 40-60% growth inhibition in contrast to the 5-10% growth inhibition of MCF7^WT^ and T47D^WT^ counterparts in response to CDK4/6 inhibitors (palbociclib -- MCF7: Fig 6E, T47D: Fig S8C and abemaciclib -- MCF7: Fig S8D, T47D: Fig S8E). These results validate in 3D spheroid assays where T47D^G55V^ cells are significantly (p=0.000004) more susceptible to palbociclib treatment than T47D^WT^ cells (Fig 6F). Lastly, *in vivo,* MCF7^G55V^ xenograft tumors regress in response to monotherapy with palbociclib, showing >95% inhibition in tumor growth compared to MCF7^WT^ tumors which show a 45% inhibition in tumor volume (Fig 6G; p=0.03, raw tumor volumes in Fig S8F: MCF7^G55V^ and Fig S8G: MCF7^WT^). Next, we tested whether susceptibility to CDK4/6 inhibitors extends to non-mutational cytoplasmic MLH1 localization by assessing *ex vivo* growth response in PDXs (Fig 6H). Both HCI003 and HCI007 (with cytoplasmic MLH1) have >50% growth inhibition after monotherapy with palbociclib (p=0.00001) while HCI017 (with nuclear MLH1) shows <10% growth inhibition (p=0.26). Similarly, *in vivo* growth in the Xenostart PDX panel indicates enrichment for palbociclib (Fig 6I) and abemaciclib (Fig 6J) responders in PDXs with cytoplasmic MLH1 relative to those with nuclear MLH1.

To test whether this causal link between cytoplasmic MLH1 and CDK4/6 sensitivity extends to patients, we investigated the correlation between sub-clonal *MLH1* mutations and treatment response in two clinical trials. In NeoPalAna, we identified two patient tumors of interest: one with *MLH1^G55V^* at 3% VAF and another with *MLH1^Q47C^*, a mutation in the same ATPase domain as G55V with 4% VAF (Fig 7A, VAFs in inset). While these tumors appear responsive to initial endocrine treatment in terms of proliferative inhibition as measured by Ki67 positivity, both tumors show significant further inhibition, 80% and 84%, respectively, in proliferation after palbociclib treatment (Fig 7B). This is in contrast to patient tumors with *MLH1^WT^,* which demonstrate 70% growth inhibition with endocrine treatment alone and little further proliferative inhibition (17%) with addition of palbociclib (Fig 7B). In a second clinical trial, NCT3007979^58^ where circulating tumor DNA was analyzed at baseline and at serial time points after cyclical CDK4/6 inhibitor treatment to evaluate circulating tumor DNA fraction and mutations in circulating tumor DNA (Fig 7C), we found an E34K mutation in the ATPase domain of *MLH1*. This mutation decreases in VAF after 3 cycles of CDK4/6 inhibition from 2.5% to undetectable (Fig 7D) unlike the VAF of *RB1* deletion, a known driver of resistance to palbociclib^59–63^, which increases after exposure to treatment from 1% to 14% (Fig 7D). In this clinical trial, patient tumors were also independently categorized as either proficient or deficient in MMR based on high blood tumor mutation burden and microsatellite instability^58^. Using this categorization, we found that the patient with *MLH1^E34K^*, as well as other patients categorized as MMR deficient show significant decrease in total circulating tumor DNA load with repeated exposure to palbociclib in contrast to patients who are proficient for MMR or have *RB1* variants, which associates with resistance to palbociclib (Fig 7E). Overall, these data suggest that sub-clonal populations bearing *MLH1* mutations clustering around the ATP-binding loop (Fig 7F-G) in the ATPase domain may seed resistance to standard endocrine therapy but prove highly susceptible to adjuvant first-line combinatorial treatment with CDK4/6 inhibitors (Fig 8).

**Figure 7:**
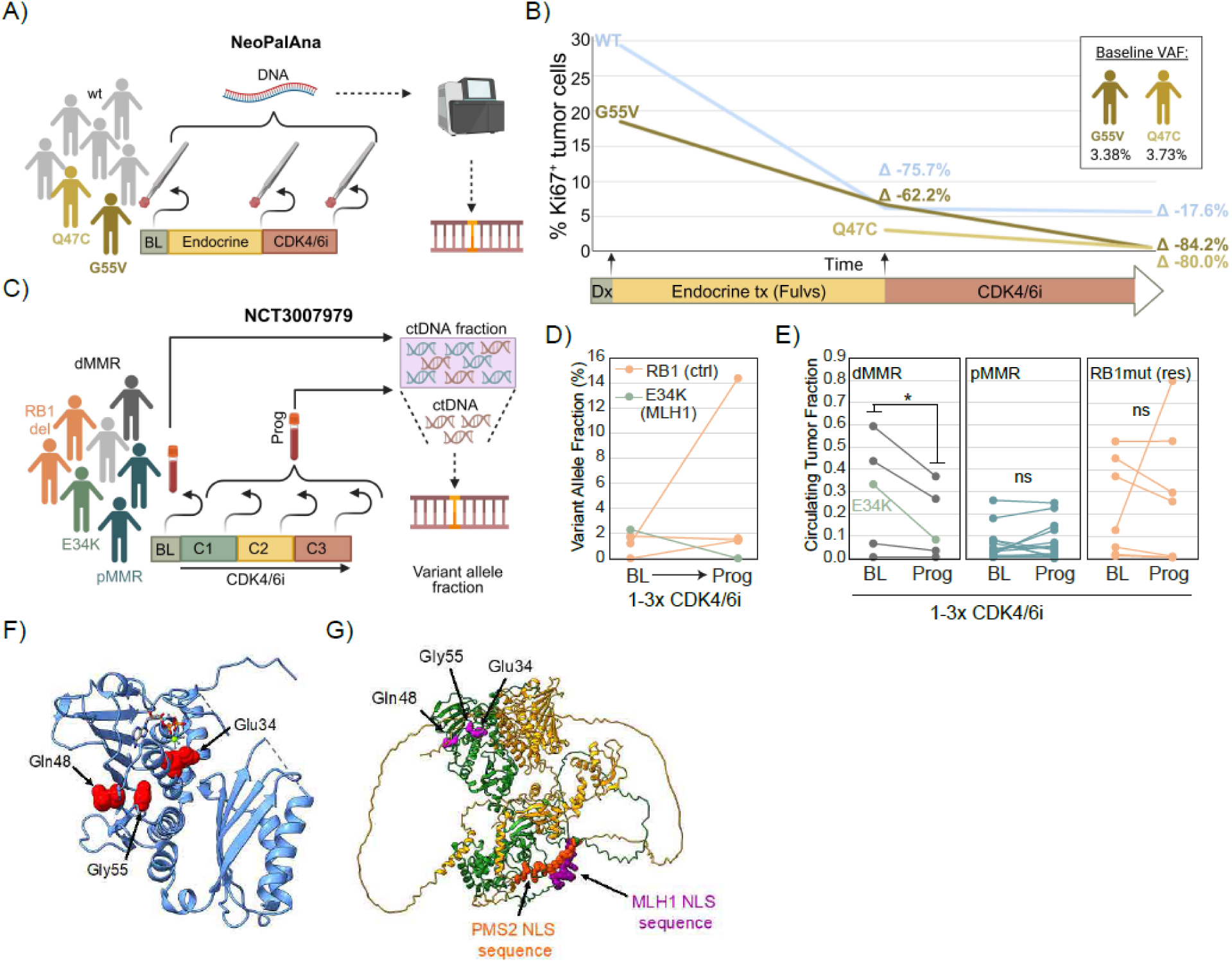
*MLH1* mutations in patient tumors and CDK4/6 inhibitor response. (A, C) Schematic of longitudinal clinical trials NeoPalAna (A) and NCT3007979 (C). (B, D-E) Line graphs showing Ki67 staining in MLH1^WT^ patient tumors compared to MLH1^G55V^ and MLH1^Q47C^ tumors after each round of treatment in NeoPalAna (B), VAF of *RB1* deletion and MLH1^E34K^ at baseline and after 3 cycles of CDK4/6 inhibitor treatment (D) and circulating tumor fraction in MMR deficient, MMR proficient and RB1-deleted patient blood samples (E) in NCT3007979. (F) Crystal structure of the N-terminal domain of human MLH1 bound to ADP (PDB: 4P7A). Residues of significance are highlighted with red spheres. (G) AlphaFold Multimer predicted structure of full-length human MLH1 (green)-PMS2 (yellow) with residues of significance shown in magenta spheres. Nuclear localization sequences are shown with purple and orange spheres for MLH1 and PMS2, respectively. Residue numbering is based on UniProt sequence and isoform number P40692-1. Student’s t-test determined all p values. *represents p≤0.05. Abbreviations: Baseline (BL), CDK4/6 inhibitor (CDKi), Cycle 1,2,3 (C1,C2,C3), Circulating DNA (ctDNA), Progression (Prog), Deficient for MMR (dMMR), Proficient for MMR (pMMR), Nuclear MLH1 (nMLH1), Cytoplasmic MLH1 (cMLH1), not significant (ns).

**Figure 8:**
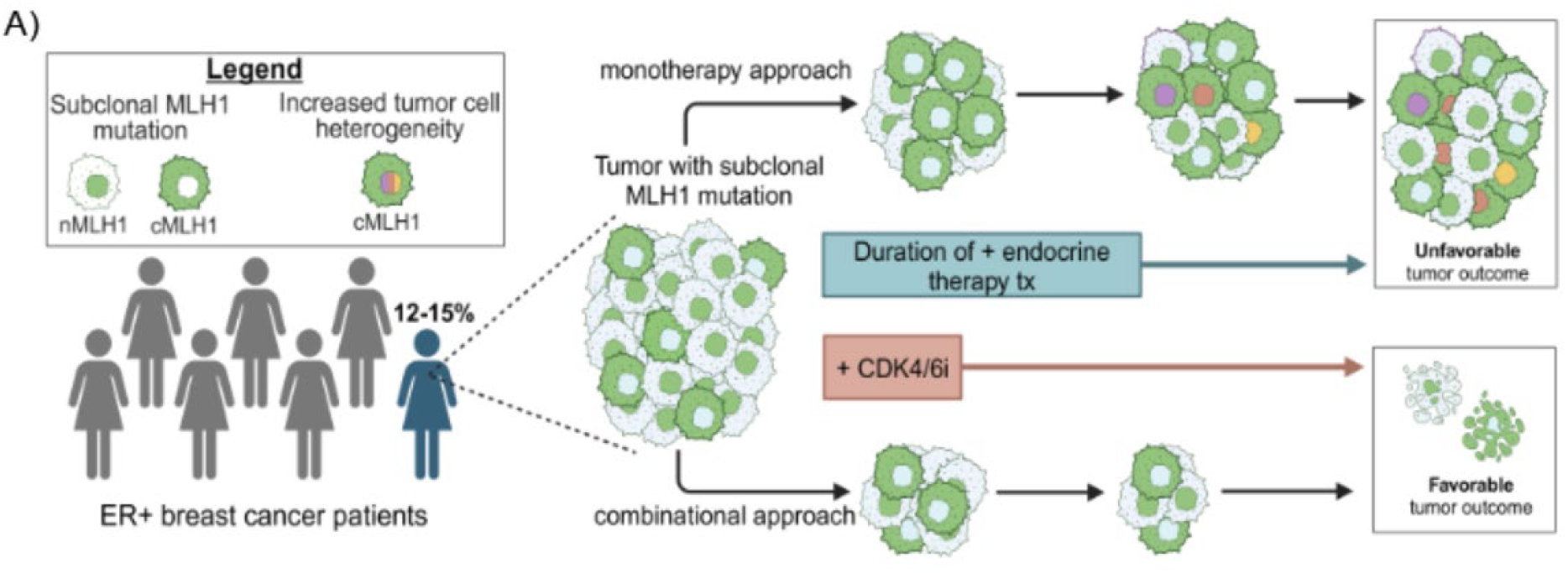
Working models: Around 12-15% of ER+/HER2- breast cancer patients have cytoplasmic MLH1 at diagnosis, in addition to the ∼12% with complete MLH1 loss. These patients may be less likely to respond to endocrine monotherapy. However, it is possible that a combinatorial approach of CDK4/6 inhibition along with endocrine therapy as first-line treatment may benefit this subset of patients.

## Discussion

The impact of our findings could be substantial given that cytoplasmic MLH1, caused by *MLH1* mutation as well as other non-mutational events, occurs in ∼12% of ER+/HER2− breast cancer patients (∼22,000 patients a year in the US alone). Currently, although MMR loss is a recognized clinical diagnostic for many cancer types^64, 65^, existing diagnostic assays either quantify presence of absence of MMR proteins using immunohistochemistry, or genomic instability by assessing tumor mutation burden or microsatellite instability^64^. However, these routine clinical diagnostics in their current iteration can neither detect sub-clonal *MLH1* mutations nor aberrant cytoplasmic localization of MLH1. Sub-clonal *MLH1* mutations are unlikely to alter genomic instability of the tumor sufficiently to be detected in assays that sequence a bulk tumor sample, and sub-cellular localization of MLH1 is not currently quantified by immunohistochemical diagnostics. However, results presented above suggest that aberrant cytoplasmic localization of MLH1 may indeed serve as a refined prognostic marker of resistance to first-line standard endocrine therapies, which along with complete absence of MLH1 could stratify 20-30% of ER+/HER2- breast cancer patients for personalized therapeutic combinations. Given the common use of MLH1 immunohistochemistry as a clinical diagnostic, it is feasible to propose the extended use of this assay to detect cytoplasmic localization of MLH1 in addition to total protein levels. Extending this assay to detect cytoplasmic localization will require additional retrospective analyses of patient tumors using the pathological grading scale presented here and assessing responsiveness to endocrine therapy in order to develop criteria for clinically relevant cytoplasmic vs nuclear characterization in the context of treatment response. Given that endocrine treatment resistance is a critical clinical problem affecting almost a third of women who are diagnosed with ER+/HER2- breast cancer across the world, such efforts would appear warranted.

The question of sub-clonal persister-like populations as seeds of endocrine treatment resistance and recurrence in ER+/HER2- breast cancer is a research area of intense focus. Previous, seminal studies have identified mutations in *ESR1*, the ER gene, and *ERBB2,* the HER2 gene as drivers of endocrine treatment resistance and disease recurrence^66–69^. In both cases, mutational fraction of these genes is low and often sub-clonal in primary ER+/HER2- breast cancer patients at diagnosis^68, 70^ but these sub-clones can grow out in the presence of endocrine treatment to eventually cause recurrence, which is almost invariably fatal^71^. The importance of identifying these mutations early on has been emphasized by the presence of existing targeted therapies (CDK4/6 inhibitors in the case of *ESR1* mutations^70^, and HER2 inhibitors in the case of *ERBB2* mutations^72^) that could prevent recurrence and breast cancer-related death for these patients. Unfortunately, in both these cases the initial VAF is so low that it is undetected, or it is posited that the mutations occur *de novo* during endocrine therapy administration^73^. Mutations in *MLH1* have a similar caveat in terms of clinical utility as they are also sub-clonal contributing to low VAF at diagnosis. In addition, mutations in *MLH1* are often novel and missense which makes determining their pathogenicity and clinical utility challenging. Assessing the functional consequences of individual *MLH1* mutations is complex due to the multifaceted nature of the MLH1 protein. With its ability to bind multiple protein partners, including itself, form complex interactions with DNA, possessing both ATPase and endonuclease activities when dimerized with a nucleolytic MutL homolog, missense mutations in *MLH1* can lead to various outcomes and impacts its many functions differently. These range from protein instability to the loss of specific functions associated with this multitude of activities^20, 74–80^.

The ATPase activity of MLH1, along with its mismatch repair heterodimer binding partners, shares an ATPase fold with histidine kinases, gyrases, and the heat shock protein 90 family^81^. The structure of the ATPase fold gives insight into how missense mutations in this domain can affect function of the protein. The ATPase fold comprises an alpha/beta sandwich formed between three alpha helices and four beta strands arranged in an antiparallel beta sheet. ATP binds between the beta sheet and an alpha helix, which is part of a longer loop structure partially enclosing ATP within the fold. The E34, G55, and Q48 residues (found to be mutated in patient tumors described above) all reside within this ATPase fold on the solved crystal structure of the human MLH1 ATPase domain bound to ADP and magnesium ion (PDB: 4P7A)^82^. Although *MLH1^G55V^* failed to localize to the nucleus, the G55V residue is not close to the MLH1 nuclear localization signal which lies in the intrinsically disordered regions connecting the ATPase domain with the dimerization interface^83, 84^. However, numerous biochemical studies of this highly conserved protein using bacterial, fungi, and mammalian models suggest that MLH1 undergoes significant conformational changes involving the intrinsically disordered region in response to ATP binding and hydrolysis in the ATPase domain^85–87^. A plausible model for the failure of ATPase domain mutated *MLH1* to transport into the nucleus is alteration or inhibition of ATPase activity causing a conformational change that hides the nuclear localization signal, thereby preventing nuclear import and dysregulating the cell cycle.

We previously showed that complete loss of MLH1 delinks cell cycle regulation in ER+ breast cancer cells from their primary growth stimulus, serum estradiol, by preventing Chk2 activation^7^. This consequently results in resistance to endocrine treatments that work by targeting the estrogen-ER signaling axis^7^. We build on this previous finding here to show that even with abundant MLH1, aberrant cytoplasmic localization of the protein significantly decreases activation of Chk2 in response to endocrine therapy. Concordantly, adding a Chk2 activator (DIM) to cells with cytoplasmic MLH1, or inhibiting downstream effectors such as CDK4/6, rescues endocrine therapy responsiveness. Many studies in the literature have identified *CHEK2* germline variants as specifically associating with increased incidence of ER+/HER2- tumors, but not other breast cancer subtypes^47, 88^. Our data further support a unique role for Chk2 in mediating the cell cycle effects of estrogen stimulus for treatment responsiveness of luminal tumors. The downstream effect of loss of Chk2 activity is dependence on G1 cyclin-dependent kinases CDK4 and CDK6 for sustaining proliferation, and therefore a therapeutic opportunity for the use of CDK4/6 inhibitors. CDK4/6 inhibitors in the metastatic setting have demonstrated significant clinical benefit in ER+/HER2- patients^42^. However, the benefit of CDK4/6 inhibition as an adjuvant treatment in early-stage ER+/HER2- patients remain conflicting^42, 89–92^. There is consensus within the cancer research community that adjuvant use of CDK4/6 inhibitors, which would greatly benefit a subset of patients, is only achievable with the identification of predictive biomarkers that can be assessed at diagnosis^41, 43, 93, 94^. Currently, there are no specific predictive biomarkers for adjuvant use of CDK4/6 inhibitors, and clinicians rely on more generic indicators of proliferation, such as high Ki67, to guide decision making on the administration of CDK4/6 inhibitors. However, the inaccuracy of Ki67 as a biomarker for administering CDK4/6 inhibitors to patients with ER+/HER2- breast cancer is widely acknowledged^95, 96^. Given that immunohistochemistry-based staining for MLH1 is routinely used in the clinic for colorectal and endometrial cancer patients^97, 98^, it may well prove feasible to administer this diagnostic for breast cancer patients along with standard immunohistochemical tests to characterize ER, PR and HER2 status. The data presented here provide rationale to set up clinical trials using immunohistochemistry-based assessment of cytoplasmic MLH1 as a companion diagnostic to test the combinatorial efficacy of CDK4/6 inhibitors and endocrine therapies as first-line treatment in a substantive subset of ER+/HER2- patients. The use of aberrant cytoplasmic localization of MLH1 along with complete loss of MLH1 as a first-in-class predictive marker for responsiveness to CDK4/6 inhibition has the potential to significantly improve breast cancer patient outcome.

## Acknowledgements

Funding: This work was supported by the Komen CCR18548157, ACS RSG-22-094-01 and R37-CA270362-02 (to S.H.); the California Institute for Regenerative Medicine, EDUC4-12813 (to A.M.); T32 CA211036-05 (to T.D) and NIGMS R35GM142651 (to C.M.M). Author contributions: Data collection, curation, and analysis: AM, TD, EO, NP, DL, MR, SE, GE, VK, IM, ME, NF, MNB, JMP, CMM, SH. Conceptual oversight: AM and SH. Preparing manuscript: AM, TD and SH. Writing and editing of manuscript: AM, TD, JMP, CMM, MNB and SH.

## References

(1) Fusco, N.; Lopez, G.; Corti, C.; Pesenti, C.; Colapietro, P.; Ercoli, G.; Gaudioso, G.; Faversani, A.; Gambini, D.; Michelotti, A.; Despini, L.; Blundo, C.; Vaira, V.; Miozzo, M.; Ferrero, S.; Bosari, S. Mismatch Repair Protein Loss as a Prognostic and Predictive Biomarker in Breast Cancers Regardless of Microsatellite Instability. JNCI Cancer Spectr. 2018, 2 (4), pky056. 10.1093/jncics/pky056.

(2) Anurag, M.; Punturi, N.; Hoog, J.; Bainbridge, M.; Ellis, M.; Haricharan, S. Comprehensive Profiling of DNA Repair Defects in Breast Cancer Identifies a Novel Class of Endocrine Therapy Resistance Drivers. Clin. Cancer Res. Off. J. Am. Assoc. Cancer Res. 2018, 24 (19), 4887–4899. 10.1158/1078-0432.CCR-17-3702.

(3) Bou Farhat, E.; Adib, E.; Daou, M.; Naqash, A. R.; Matulonis, U.; Ng, K.; Kwiatkowski, D. J.; Sholl, L. M.; Nassar, A. H. Benchmarking Mismatch Repair Testing for Patients with Cancer Receiving Immunotherapy. Cancer Cell 2024, 42 (1), 6–7. 10.1016/j.ccell.2023.12.001.

(4) Vilar, E.; Gruber, S. B. Microsatellite Instability in Colorectal Cancer—the Stable Evidence. Nat. Rev. Clin. Oncol. 2010, 7 (3), 153–162. 10.1038/nrclinonc.2009.237.

(5) Latham, A.; Srinivasan, P.; Kemel, Y.; Shia, J.; Bandlamudi, C.; Mandelker, D.; Middha, S.; Hechtman, J.; Zehir, A.; Dubard-Gault, M.; Tran, C.; Stewart, C.; Sheehan, M.; Penson, A.; DeLair, D.; Yaeger, R.; Vijai, J.; Mukherjee, S.; Galle, J.; Dickson, M. A.; Janjigian, Y.; O’Reilly, E. M.; Segal, N.; Saltz, L. B.; Reidy-Lagunes, D.; Varghese, A. M.; Bajorin, D.; Carlo, M. I.; Cadoo, K.; Walsh, M. F.; Weiser, M.; Aguilar, J. G.; Klimstra, D. S.; Diaz, L. A.; Baselga, J.; Zhang, L.; Ladanyi, M.; Hyman, D. M.; Solit, D. B.; Robson, M. E.; Taylor, B. S.; Offit, K.; Berger, M. F.; Stadler, Z. K. Microsatellite Instability Is Associated With the Presence of Lynch Syndrome Pan-Cancer. J. Clin. Oncol. 2019, 37 (4), 286–295. 10.1200/JCO.18.00283.

(6) Berg, H. F.; Engerud, H.; Myrvold, M.; Lien, H. E.; Hjelmeland, M. E.; Halle, M. K.; Woie, K.; Hoivik, E. A.; Haldorsen, I. S.; Vintermyr, O.; Trovik, J.; Krakstad, C. Mismatch Repair Markers in Preoperative and Operative Endometrial Cancer Samples; Expression Concordance and Prognostic Value. Br. J. Cancer 2023, 128 (4), 647–655. 10.1038/s41416-022-02063-3.

(7) Haricharan, S.; Punturi, N.; Singh, P.; Holloway, K. R.; Anurag, M.; Schmelz, J.; Schmidt, C.; Lei, J. T.; Suman, V.; Hunt, K.; Olson, J. A.; Hoog, J.; Li, S.; Huang, S.; Edwards, D. P.; Kavuri, S. M.; Bainbridge, M. N.; Ma, C. X.; Ellis, M. J. Loss of MutL Disrupts CHK2-Dependent Cell-Cycle Control through CDK4/6 to Promote Intrinsic Endocrine Therapy Resistance in Primary Breast Cancer. Cancer Discov. 2017, 7 (10), 1168–1183. 10.1158/2159-8290.CD-16-1179.

(8) Kane, M. F.; Loda, M.; Gaida, G. M.; Lipman, J.; Mishra, R.; Goldman, H.; Jessup, J. M.; Kolodner, R. Methylation of the HMLH1 Promoter Correlates with Lack of Expression of HMLH1 in Sporadic Colon Tumors and Mismatch Repair-Defective Human Tumor Cell Lines. Cancer Res. 1997, 57 (5), 808–811.

(9) Muquith, M.; Espinoza, M.; Elliott, A.; Xiu, J.; Seeber, A.; El-Deiry, W.; Antonarakis, E. S.; Graff, S. L.; Hall, M. J.; Borghaei, H.; Hoon, D. S. B.; Liu, S. V.; Ma, P. C.; McKay, R. R.; Wise-Draper, T.; Marshall, J.; Sledge, G. W.; Spetzler, D.; Zhu, H.; Hsiehchen, D. Tissue-Specific Thresholds of Mutation Burden Associated with Anti-PD-1/L1 Therapy Benefit and Prognosis in Microsatellite-Stable Cancers. *Nat*. Cancer 2024, 1–9. 10.1038/s43018-024-00752-x.

(10) Sajjadi, E.; Venetis, K.; Piciotti, R.; Invernizzi, M.; Guerini-Rocco, E.; Haricharan, S.; Fusco, N. Mismatch Repair-Deficient Hormone Receptor-Positive Breast Cancers: Biology and Pathological Characterization. Cancer Cell Int. 2021, 21 (1), 266. 10.1186/s12935-021-01976-y.

(11) Donehower, L. A.; Creighton, C. J.; Schultz, N.; Shinbrot, E.; Chang, K.; Gunaratne, P. H.; Muzny, D.; Sander, C.; Hamilton, S. R.; Gibbs, R. A.; Wheeler, D. MLH1-Silenced and Non-Silenced Subgroups of Hypermutated Colorectal Carcinomas Have Distinct Mutational Landscapes. J. Pathol. 2013, 229 (1), 99–110. 10.1002/path.4087.

(12) Lei, J. T.; Anurag, M.; Haricharan, S.; Gou, X.; Ellis, M. J. Endocrine Therapy Resistance: New Insights. Breast Edinb. Scotl. 2019, 48 Suppl 1 (Suppl 1), S26–S30. 10.1016/S0960-9776(19)31118-X.

(13) Punturi, N. B.; Seker, S.; Devarakonda, V.; Mazumder, A.; Kalra, R.; Chen, C. H.; Li, S.; Primeau, T.; Ellis, M. J.; Kavuri, S. M.; Haricharan, S. Mismatch Repair Deficiency Predicts Response to HER2 Blockade in HER2-Negative Breast Cancer. Nat. Commun. 2021, 12 (1), 2940. 10.1038/s41467-021-23271-0.

(14) Zukin, E.; Culver, J. O.; Liu, Y.; Yang, Y.; Ricker, C. N.; Hodan, R.; Sturgeon, D.; Kingham, K.; Chun, N. M.; Rowe-Teeter, C.; Singh, K.; Zell, J. A.; Ladabaum, U.; McDonnell, K. J.; Ford, J. M.; Parmigiani, G.; Braun, D.; Kurian, A. W.; Gruber, S. B.; Idos, G. E. Clinical Implications of Conflicting Variant Interpretations in the Cancer Genetics Clinic. Genet. Med. Off. J. Am. Coll. Med. Genet. 2023, 25 (7), 100837. 10.1016/j.gim.2023.100837.

(15) Drost, M.; Tiersma, Y.; Thompson, B. A.; Frederiksen, J. H.; Keijzers, G.; Glubb, D.; Kathe, S.; Osinga, J.; Westers, H.; Pappas, L.; Boucher, K. M.; Molenkamp, S.; Zonneveld, J. B.; van Asperen, C. J.; Goldgar, D. E.; Wallace, S. S.; Sijmons, R. H.; Spurdle, A. B.; Rasmussen, L. J.; Greenblatt, M. S.; de Wind, N.; Tavtigian, S. V. A Functional Assay-Based Procedure to Classify Mismatch Repair Gene Variants in Lynch Syndrome. Genet. Med. Off. J. Am. Coll. Med. Genet. 2019, 21 (7), 1486–1496. 10.1038/s41436-018-0372-2.

(16) Köger, N.; Paulsen, L.; López-Kostner, F.; Della Valle, A.; Vaccaro, C. A.; Palmero, E. I.; Alvarez, K.; Sarroca, C.; Neffa, F.; Kalfayan, P. G.; Gonzalez, M. L.; Rossi, B. M.; Reis, R. M.; Brieger, A.; Zeuzem, S.; Hinrichsen, I.; Dominguez-Valentin, M.; Plotz, G. Evaluation of MLH1 Variants of Unclear Significance. Genes. Chromosomes Cancer 2018, 57 (7), 350– 358. 10.1002/gcc.22536.

(17) Rath, A.; Radecki, A. A.; Rahman, K.; Gilmore, R. B.; Hudson, J. R.; Cenci, M.; Tavtigian, S. V.; Grady, J. P.; Heinen, C. D. A Calibrated Cell-Based Functional Assay to Aid Classification of MLH1 DNA Mismatch Repair Gene Variants. Hum. Mutat. 2022, 43 (12), 2295–2307. 10.1002/humu.24462.

(18) Martinez, S. L.; Kolodner, R. D. Functional Analysis of Human Mismatch Repair Gene Mutations Identifies Weak Alleles and Polymorphisms Capable of Polygenic Interactions. Proc. Natl. Acad. Sci. U. S. A. 2010, 107 (11), 5070–5075. 10.1073/pnas.1000798107.

(19) Shimodaira, H.; Filosi, N.; Shibata, H.; Suzuki, T.; Radice, P.; Kanamaru, R.; Friend, S. H.; Kolodner, R. D.; Ishioka, C. Functional Analysis of Human MLH1 Mutations in Saccharomyces Cerevisiae. Nat. Genet. 1998, 19 (4), 384–389. 10.1038/1277.

(20) Manhart, C. M.; Ni, X.; White, M. A.; Ortega, J.; Surtees, J. A.; Alani, E. The Mismatch Repair and Meiotic Recombination Endonuclease Mlh1-Mlh3 Is Activated by Polymer Formation and Can Cleave DNA Substrates in Trans. PLoS Biol. 2017, 15 (4), e2001164. 10.1371/journal.pbio.2001164.

(21) Luo, Y.; Lin, F.-T.; Lin, W.-C. ATM-Mediated Stabilization of HMutL DNA Mismatch Repair Proteins Augments P53 Activation during DNA Damage. Mol. Cell. Biol. 2004, 24 (14), 6430–6444. 10.1128/MCB.24.14.6430-6444.2004.

(22) Gibson, S. L.; Bindra, R. S.; Glazer, P. M. Hypoxia-Induced Phosphorylation of Chk2 in an Ataxia Telangiectasia Mutated-Dependent Manner. Cancer Res. 2005, 65 (23), 10734– 10741. 10.1158/0008-5472.CAN-05-1160.

(23) Abildgaard, A. B.; Nielsen, S. V.; Bernstein, I.; Stein, A.; Lindorff-Larsen, K.; Hartmann-Petersen, R. Lynch Syndrome, Molecular Mechanisms and Variant Classification. Br. J. Cancer 2023, 128 (5), 726–734. 10.1038/s41416-022-02059-z.

(24) Chen, F.; Ding, K.; Priedigkeit, N.; Elangovan, A.; Levine, K. M.; Carleton, N.; Savariau, L.; Atkinson, J. M.; Oesterreich, S.; Lee, A. V. Single-Cell Transcriptomic Heterogeneity in Invasive Ductal and Lobular Breast Cancer Cells. Cancer Res. 2021, 81 (2), 268–281. 10.1158/0008-5472.CAN-20-0696.

(25) McGranahan, N.; Swanton, C. Clonal Heterogeneity and Tumor Evolution: Past, Present, and the Future. Cell 2017, 168 (4), 613–628. 10.1016/j.cell.2017.01.018.

(26) Pribluda, A.; de la Cruz, C. C.; Jackson, E. L. Intratumoral Heterogeneity: From Diversity Comes Resistance. Clin. Cancer Res. Off. J. Am. Assoc. Cancer Res. 2015, 21 (13), 2916–2923. 10.1158/1078-0432.CCR-14-1213.

(27) Pu, Y.; Li, L.; Peng, H.; Liu, L.; Heymann, D.; Robert, C.; Vallette, F.; Shen, S. Drug-Tolerant Persister Cells in Cancer: The Cutting Edges and Future Directions. Nat. Rev. Clin. Oncol. 2023, 20 (11), 799–813. 10.1038/s41571-023-00815-5.

(28) Dhanyamraju, P. K.; Schell, T. D.; Amin, S.; Robertson, G. P. Drug-Tolerant Persister Cells in Cancer Therapy Resistance. Cancer Res. 2022, 82 (14), 2503–2514. 10.1158/0008-5472.CAN-21-3844.

(29) Haricharan, S.; Bainbridge, M. N.; Scheet, P.; Brown, P. H. Somatic Mutation Load of Estrogen Receptor-Positive Breast Tumors Predicts Overall Survival: An Analysis of Genome Sequence Data. Breast Cancer Res. Treat. 2014, 146 (1), 211–220. 10.1007/s10549-014-2991-x.

(30) Fittall, M. W.; Van Loo, P. Translating Insights into Tumor Evolution to Clinical Practice: Promises and Challenges. Genome Med. 2019, 11, 20. 10.1186/s13073-019-0632-z.

(31) Yates, L. R.; Knappskog, S.; Wedge, D.; Farmery, J. H. R.; Gonzalez, S.; Martincorena, I.; Alexandrov, L. B.; Van Loo, P.; Haugland, H. K.; Lilleng, P. K.; Gundem, G.; Gerstung, M.; Pappaemmanuil, E.; Gazinska, P.; Bhosle, S. G.; Jones, D.; Raine, K.; Mudie, L.; Latimer, C.; Sawyer, E.; Desmedt, C.; Sotiriou, C.; Stratton, M. R.; Sieuwerts, A. M.; Lynch, A. G.; Martens, J. W.; Richardson, A. L.; Tutt, A.; Lønning, P. E.; Campbell, P. J. Genomic Evolution of Breast Cancer Metastasis and Relapse. Cancer Cell 2017, 32 (2), 169–184.e7. 10.1016/j.ccell.2017.07.005.

(32) Hanker, A. B.; Sudhan, D. R.; Arteaga, C. L. Overcoming Endocrine Resistance in Breast Cancer. Cancer Cell 2020, 37 (4), 496–513. 10.1016/j.ccell.2020.03.009.

(33) Razavi, P.; Chang, M. T.; Xu, G.; Bandlamudi, C.; Ross, D. S.; Vasan, N.; Cai, Y.; Bielski, C. M.; Donoghue, M. T. A.; Jonsson, P.; Penson, A.; Shen, R.; Pareja, F.; Kundra, R.; Middha, S.; Cheng, M. L.; Zehir, A.; Kandoth, C.; Patel, R.; Huberman, K.; Smyth, L. M.; Jhaveri, K.; Modi, S.; Traina, T. A.; Dang, C.; Zhang, W.; Weigelt, B.; Li, B. T.; Ladanyi, M.; Hyman, D. M.; Schultz, N.; Robson, M. E.; Hudis, C.; Brogi, E.; Viale, A.; Norton, L.; Dickler, M. N.; Berger, M. F.; Iacobuzio-Donahue, C. A.; Chandarlapaty, S.; Scaltriti, M.; Reis-Filho, J. S.; Solit, D. B.; Taylor, B. S.; Baselga, J. The Genomic Landscape of Endocrine-Resistant Advanced Breast Cancers. Cancer Cell 2018, 34 (3), 427–438.e6. 10.1016/j.ccell.2018.08.008.

(34) Shao, W.; Brown, M. Advances in Estrogen Receptor Biology: Prospects for Improvements in Targeted Breast Cancer Therapy. Breast Cancer Res. BCR 2004, 6 (1), 39–52. 10.1186/bcr742.

(35) Venetis, K.; Sajjadi, E.; Haricharan, S.; Fusco, N. Mismatch Repair Testing in Breast Cancer: The Path to Tumor-Specific Immuno-Oncology Biomarkers. Transl. Cancer Res. 2020, 9 (7), 4060–4064. 10.21037/tcr-20-1852.

(36) Ellis, M. Overcoming Endocrine Therapy Resistance by Signal Transduction Inhibition. The Oncologist 2004, 9 *Suppl 3*, 20–26. 10.1634/theoncologist.9-suppl_3-20.

(37) O’Leary, B.; Finn, R. S.; Turner, N. C. Treating Cancer with Selective CDK4/6 Inhibitors. Nat. Rev. Clin. Oncol. 2016, 13 (7), 417–430. 10.1038/nrclinonc.2016.26.

(38) Awada, A.; Gligorov, J.; Jerusalem, G.; Preusser, M.; Singer, C.; Zielinski, C. CDK4/6 Inhibition in Low Burden and Extensive Metastatic Breast Cancer: Summary of an *ESMO Open—Cancer Horizons* pro and Con Discussion. ESMO Open 2019, 4 (6), e000565. 10.1136/esmoopen-2019-000565.

(39) Turner Nicholas C.; Slamon Dennis J.; Ro Jungsil; Bondarenko Igor; Im Seock-Ah; Masuda Norikazu; Colleoni Marco; DeMichele Angela; Loi Sherene; Verma Sunil; Iwata Hiroji; Harbeck Nadia; Loibl Sibylle; André Fabrice; Puyana Theall Kathy; Huang Xin; Giorgetti Carla; Huang Bartlett Cynthia; Cristofanilli Massimo. Overall Survival with Palbociclib and Fulvestrant in Advanced Breast Cancer. N. Engl. J. Med. 2018, 379 (20), 1926–1936. 10.1056/NEJMoa1810527.

(40) Moore, D. C.; Guinigundo, A. S. Revolutionizing Cancer Treatment: Harnessing the Power of Biomarkers to Improve Patient Outcomes. J. Adv. Pract. Oncol. 2023, 14 (Suppl 1), 4–8. 10.6004/jadpro.2023.14.3.15.

(41) Anurag, M.; Haricharan, S.; Ellis, M. J. CDK4/6 Inhibitor Biomarker Research: Are We Barking Up the Wrong Tree? Clin. Cancer Res. Off. J. Am. Assoc. Cancer Res. 2020, 26 (1), 3–5. 10.1158/1078-0432.CCR-19-3119.

(42) Morrison, L.; Loibl, S.; Turner, N. C. The CDK4/6 Inhibitor Revolution — a Game-Changing Era for Breast Cancer Treatment. Nat. Rev. Clin. Oncol. 2024, 21 (2), 89–105. 10.1038/s41571-023-00840-4.

(43) Schoninger, S. F.; Blain, S. W. The Ongoing Search for Biomarkers of CDK4/6 Inhibitor Responsiveness in Breast Cancer. Mol. Cancer Ther. 2020, 19 (1), 3–12. 10.1158/1535-7163.MCT-19-0253.

(44) Migliaccio, I.; Bonechi, M.; McCartney, A.; Guarducci, C.; Benelli, M.; Biganzoli, L.; Di Leo, A.; Malorni, L. CDK4/6 Inhibitors: A Focus on Biomarkers of Response and Post-Treatment Therapeutic Strategies in Hormone Receptor-Positive HER2-Negative Breast Cancer. Cancer Treat. Rev. 2021, 93, 102136. 10.1016/j.ctrv.2020.102136.

(45) Smith, J.; Tho, L. M.; Xu, N.; Gillespie, D. A. The ATM-Chk2 and ATR-Chk1 Pathways in DNA Damage Signaling and Cancer. Adv. Cancer Res. 2010, 108, 73–112. 10.1016/B978-0-12-380888-2.00003-0.

(46) Feitelson, M. A.; Arzumanyan, A.; Kulathinal, R. J.; Blain, S. W.; Holcombe, R. F.; Mahajna, J.; Marino, M.; Martinez-Chantar, M. L.; Nawroth, R.; Sanchez-Garcia, I.; Sharma, D.; Saxena, N. K.; Singh, N.; Vlachostergios, P. J.; Guo, S.; Honoki, K.; Fujii, H.; Georgakilas, A. G.; Amedei, A.; Niccolai, E.; Amin, A.; Ashraf, S. S.; Boosani, C. S.; Guha, G.; Ciriolo, M. R.; Aquilano, K.; Chen, S.; Mohammed, S. I.; Azmi, A. S.; Bhakta, D.; Halicka, D.; Nowsheen, S. Sustained Proliferation in Cancer: Mechanisms and Novel Therapeutic Targets. Semin. Cancer Biol. 2015, 35 (Suppl), S25–S54. 10.1016/j.semcancer.2015.02.006.

(47) Oropeza, E.; Seker, S.; Carrel, S.; Mazumder, A.; Lozano, D.; Jimenez, A.; VandenHeuvel, S. N.; Noltensmeyer, D. A.; Punturi, N. B.; Lei, J. T.; Lim, B.; Waltz, S. E.; Raghavan, S. A.; Bainbridge, M. N.; Haricharan, S. Molecular Portraits of Cell Cycle Checkpoint Kinases in Cancer Evolution, Progression, and Treatment Responsiveness. Sci. Adv. 2023, 9 (26), eadf2860. 10.1126/sciadv.adf2860.

(48) Toss, A.; Tenedini, E.; Piombino, C.; Venturelli, M.; Marchi, I.; Gasparini, E.; Barbieri, E.; Razzaboni, E.; Domati, F.; Caggia, F.; Grandi, G.; Combi, F.; Tazzioli, G.; Dominici, M.; Tagliafico, E.; Cortesi, L. Clinicopathologic Profile of Breast Cancer in Germline ATM and CHEK2 Mutation Carriers. Genes 2021, 12 (5), 616. 10.3390/genes12050616.

(49) Cybulski, C.; Wokołorczyk, D.; Jakubowska, A.; Huzarski, T.; Byrski, T.; Gronwald, J.; Masojć, B.; Deebniak, T.; Górski, B.; Blecharz, P.; Narod, S. A.; Lubiński, J. Risk of Breast Cancer in Women with a CHEK2 Mutation with and without a Family History of Breast Cancer. J. Clin. Oncol. Off. J. Am. Soc. Clin. Oncol. 2011, 29 (28), 3747–3752. 10.1200/JCO.2010.34.0778.

(50) Breast Cancer Association Consortium; Mavaddat, N.; Dorling, L.; Carvalho, S.; Allen, J.; González-Neira, A.; Keeman, R.; Bolla, M. K.; Dennis, J.; Wang, Q.; Ahearn, T. U.; Andrulis, I. L.; Beckmann, M. W.; Behrens, S.; Benitez, J.; Bermisheva, M.; Blomqvist, C.; Bogdanova, N. V.; Bojesen, S. E.; Briceno, I.; Brüning, T.; Camp, N. J.; Campbell, A.; Castelao, J. E.; Chang-Claude, J.; Chanock, S. J.; Chenevix-Trench, G.; Christiansen, H.; Czene, K.; Dörk, T.; Eriksson, M.; Evans, D. G.; Fasching, P. A.; Figueroa, J. D.; Flyger, H.; Gabrielson, M.; Gago-Dominguez, M.; Geisler, J.; Giles, G. G.; Guénel, P.; Hadjisavvas, A.; Hahnen, E.; Hall, P.; Hamann, U.; Hartikainen, J. M.; Hartman, M.; Hoppe, R.; Howell, A.; Jakubowska, A.; Jung, A.; Khusnutdinova, E. K.; Kristensen, V. N.; Li, J.; Lim, S. H.; Lindblom, A.; Loizidou, M. A.; Lophatananon, A.; Lubinski, J.; Madsen, M. J.; Mannermaa, A.; Manoochehri, M.; Margolin, S.; Mavroudis, D.; Milne, R. L.; Mohd Taib, N. A.; Morra, A.; Muir, K.; Obi, N.; Osorio, A.; Park-Simon, T.-W.; Peterlongo, P.; Radice, P.; Saloustros, E.; Sawyer, E. J.; Schmutzler, R. K.; Shah, M.; Sim, X.; Southey, M. C.; Thorne, H.; Tomlinson, I.; Torres, D.; Truong, T.; Yip, C. H.; Spurdle, A. B.; Vreeswijk, M. P. G.; Dunning, A. M.; García-Closas, M.; Pharoah, P. D. P.; Kvist, A.; Muranen, T. A.; Nevanlinna, H.; Teo, S. H.; Devilee, P.; Schmidt, M. K.; Easton, D. F. Pathology of Tumors Associated With Pathogenic Germline Variants in 9 Breast Cancer Susceptibility Genes. JAMA Oncol. 2022, 8 (3), e216744. 10.1001/jamaoncol.2021.6744.

(51) Loveday, C.; Garrett, A.; Law, P.; Hanks, S.; Poyastro-Pearson, E.; Adlard, J. W.; Barwell, J.; Berg, J.; Brady, A. F.; Brewer, C.; Chapman, C.; Cook, J.; Davidson, R.; Donaldson, A.; Douglas, F.; Greenhalgh, L.; Henderson, A.; Izatt, L.; Kumar, A.; Lalloo, F.; Miedzybrodzka, Z.; Morrison, P. J.; Paterson, J.; Porteous, M.; Rogers, M. T.; Walker, L.; Breast and Ovarian Cancer Susceptibility Collaboration; Eccles, D.; Evans, D. G.; Snape, K.; Hanson, H.; Houlston, R. S.; Turnbull, C. Analysis of Rare Disruptive Germline Mutations in 2135 Enriched BRCA-Negative Breast Cancers Excludes Additional High-Impact Susceptibility Genes. Ann. Oncol. Off. J. Eur. Soc. Med. Oncol. 2022, 33 (12), 1318–1327. 10.1016/j.annonc.2022.09.152.

(52) Benachenhou, N.; Guiral, S.; Gorska-Flipot, I.; Labuda, D.; Sinnett, D. Frequent Loss of Heterozygosity at the DNA Mismatch-Repair Loci HMLH1 and HMSH3 in Sporadic Breast Cancer. Br. J. Cancer 1999, 79 (7–8), 1012–1017. 10.1038/sj.bjc.6690162.

(53) Malik, S. S.; Masood, N.; Asif, M.; Ahmed, P.; Shah, Z. U.; Khan, J. S. Expressional Analysis of MLH1 and MSH2 in Breast Cancer. Curr. Probl. Cancer 2019, 43 (2), 97–105. 10.1016/j.currproblcancer.2018.08.001.

(54) Aliouat-Denis, C.-M.; Dendouga, N.; Van den Wyngaert, I.; Goehlmann, H.; Steller, U.; van de Weyer, I.; Van Slycken, N.; Andries, L.; Kass, S.; Luyten, W.; Janicot, M.; Vialard, J. E. P53-Independent Regulation of P21Waf1/Cip1 Expression and Senescence by Chk2. Mol. Cancer Res. MCR 2005, 3 (11), 627–634. 10.1158/1541-7786.MCR-05-0121.

(55) Mazumder, A.; Shiao, S.; Haricharan, S. HER2 Activation and Endocrine Treatment Resistance in HER2-Negative Breast Cancer. Endocrinology 2021, 162 (10), bqab153. 10.1210/endocr/bqab153.

(56) Brown, K. D.; Rathi, A.; Kamath, R.; Beardsley, D. I.; Zhan, Q.; Mannino, J. L.; Baskaran, R. The Mismatch Repair System Is Required for S-Phase Checkpoint Activation. Nat. Genet. 2003, 33 (1), 80–84. 10.1038/ng1052.

(57) Akli, S.; Zheng, P.-J.; Multani, A. S.; Wingate, H. F.; Pathak, S.; Zhang, N.; Tucker, S. L.; Chang, S.; Keyomarsi, K. Tumor-Specific Low Molecular Weight Forms of Cyclin E Induce Genomic Instability and Resistance to P21, P27, and Antiestrogens in Breast Cancer. Cancer Res. 2004, 64 (9), 3198–3208. 10.1158/0008-5472.can-03-3672.

(58) Krishnamurthy, J.; Luo, J.; Suresh, R.; Ademuyiwa, F.; Rigden, C.; Rearden, T.; Clifton, K.; Weilbaecher, K.; Frith, A.; Roshal, A.; Tandra, P. K.; Cherian, M.; Summa, T.; Haas, B.; Thomas, S.; Hernandez-Aya, L.; Bergqvist, M.; Peterson, L.; Ma, C. X. A Phase II Trial of an Alternative Schedule of Palbociclib and Embedded Serum TK1 Analysis. NPJ Breast Cancer 2022, 8 (1), 35. 10.1038/s41523-022-00399-w.

(59) Palafox, M.; Monserrat, L.; Bellet, M.; Villacampa, G.; Gonzalez-Perez, A.; Oliveira, M.; Brasó-Maristany, F.; Ibrahimi, N.; Kannan, S.; Mina, L.; Herrera-Abreu, M. T.; Òdena, A.; Sánchez-Guixé, M.; Capelán, M.; Azaro, A.; Bruna, A.; Rodríguez, O.; Guzmán, M.; Grueso, J.; Viaplana, C.; Hernández, J.; Su, F.; Lin, K.; Clarke, R. B.; Caldas, C.; Arribas, J.; Michiels, S.; García-Sanz, A.; Turner, N. C.; Prat, A.; Nuciforo, P.; Dienstmann, R.; Verma, C. S.; Lopez-Bigas, N.; Scaltriti, M.; Arnedos, M.; Saura, C.; Serra, V. High P16 Expression and Heterozygous RB1 Loss Are Biomarkers for CDK4/6 Inhibitor Resistance in ER+ Breast Cancer. Nat. Commun. 2022, 13 (1), 5258. 10.1038/s41467-022-32828-6.

(60) Antonarelli, G.; Taurelli Salimbeni, B.; Marra, A.; Esposito, A.; Locatelli, M. A.; Trapani, D.; Pescia, C.; Fusco, N.; Curigliano, G.; Criscitiello, C. The CDK4/6 Inhibitors Biomarker Landscape: The Most Relevant Biomarkers of Response or Resistance for Further Research and Potential Clinical Utility. Crit. Rev. Oncol. Hematol. 2023, 192, 104148. 10.1016/j.critrevonc.2023.104148.

(61) Herrera-Abreu, M. T.; Palafox, M.; Asghar, U.; Rivas, M. A.; Cutts, R. J.; Garcia-Murillas, I.; Pearson, A.; Guzman, M.; Rodriguez, O.; Grueso, J.; Bellet, M.; Cortés, J.; Elliott, R.; Pancholi, S.; Baselga, J.; Dowsett, M.; Martin, L.-A.; Turner, N. C.; Serra, V. Early Adaptation and Acquired Resistance to CDK4/6 Inhibition in Estrogen Receptor-Positive Breast Cancer. Cancer Res. 2016, 76 (8), 2301–2313. 10.1158/0008-5472.CAN-15-0728.

(62) Knudsen, E. S.; Pruitt, S. C.; Hershberger, P. A.; Witkiewicz, A. K.; Goodrich, D. W. Cell Cycle and Beyond: Exploiting New RB1 Controlled Mechanisms for Cancer Therapy. Trends Cancer 2019, 5 (5), 308–324. 10.1016/j.trecan.2019.03.005.

(63) O’Leary, B.; Cutts, R. J.; Liu, Y.; Hrebien, S.; Huang, X.; Fenwick, K.; André, F.; Loibl, S.; Loi, S.; Garcia-Murillas, I.; Cristofanilli, M.; Huang Bartlett, C.; Turner, N. C. The Genetic Landscape and Clonal Evolution of Breast Cancer Resistance to Palbociclib plus Fulvestrant in the PALOMA-3 Trial. Cancer Discov. 2018, 8 (11), 1390–1403. 10.1158/2159-8290.CD-18-0264.

(64) Buza, N.; Ziai, J.; Hui, P. Mismatch Repair Deficiency Testing in Clinical Practice. Expert Rev. Mol. Diagn. 2016, 16 (5), 591–604. 10.1586/14737159.2016.1156533.

(65) Stelloo, E.; Jansen, A. M. L.; Osse, E. M.; Nout, R. A.; Creutzberg, C. L.; Ruano, D.; Church, D. N.; Morreau, H.; Smit, V. T. H. B. M.; Wezel, T. van; Bosse, T. Practical Guidance for Mismatch Repair-Deficiency Testing in Endometrial Cancer. Ann. Oncol. 2017, 28 (1), 96–102. 10.1093/annonc/mdw542.

(66) Jeselsohn, R.; Buchwalter, G.; De Angelis, C.; Brown, M.; Schiff, R. ESR1 Mutations as a Mechanism for Acquired Endocrine Resistance in Breast Cancer. Nat. Rev. Clin. Oncol. 2015, 12 (10), 573–583. 10.1038/nrclinonc.2015.117.

(67) Fuqua, S. A. W.; Gu, G.; Rechoum, Y. Estrogen Receptor (ER) α Mutations in Breast Cancer: Hidden in Plain Sight. Breast Cancer Res. Treat. 2014, 144 (1), 11–19. 10.1007/s10549-014-2847-4.

(68) Li, S.; Shen, D.; Shao, J.; Crowder, R.; Liu, W.; Prat, A.; He, X.; Liu, S.; Hoog, J.; Lu, C.; Ding, L.; Griffith, O. L.; Miller, C.; Larson, D.; Fulton, R. S.; Harrison, M.; Mooney, T.; McMichael, J. F.; Luo, J.; Tao, Y.; Goncalves, R.; Schlosberg, C.; Hiken, J. F.; Saied, L.; Sanchez, C.; Giuntoli, T.; Bumb, C.; Cooper, C.; Kitchens, R. T.; Lin, A.; Phommaly, C.; Davies, S. R.; Zhang, J.; Kavuri, M. S.; McEachern, D.; Dong, Y. Y.; Ma, C.; Pluard, T.; Naughton, M.; Bose, R.; Suresh, R.; McDowell, R.; Michel, L.; Aft, R.; Gillanders, W.; DeSchryver, K.; Wilson, R. K.; Wang, S.; Mills, G. B.; Gonzalez-Angulo, A.; Edwards, J. R.; Maher, C.; Perou, C. M.; Mardis, E. R.; Ellis, M. J. Endocrine-Therapy-Resistant ESR1 Variants Revealed by Genomic Characterization of Breast-Cancer-Derived Xenografts. Cell Rep. 2013, 4 (6), 10.1016/j.celrep.2013.08.022. 10.1016/j.celrep.2013.08.022.

(69) Bose, R.; Kavuri, S. M.; Searleman, A. C.; Shen, W.; Shen, D.; Koboldt, D. C.; Monsey, J.; Goel, N.; Aronson, A. B.; Li, S.; Ma, C. X.; Ding, L.; Mardis, E. R.; Ellis, M. J. Activating HER2 Mutations in HER2 Gene Amplification Negative Breast Cancer. Cancer Discov. 2013, 3 (2), 224–237. 10.1158/2159-8290.CD-12-0349.

(70) Brett, J. O.; Spring, L. M.; Bardia, A.; Wander, S. A. ESR1 Mutation as an Emerging Clinical Biomarker in Metastatic Hormone Receptor-Positive Breast Cancer. Breast Cancer Res. 2021, 23 (1), 85. 10.1186/s13058-021-01462-3.

(71) Mavrommati, I.; Johnson, F.; Echeverria, G. V.; Natrajan, R. Subclonal Heterogeneity and Evolution in Breast Cancer. Npj Breast Cancer 2021, 7 (1), 1–9. 10.1038/s41523-021-00363-0.

(72) Yi, Z.; Rong, G.; Guan, Y.; Li, J.; Chang, L.; Li, H.; Liu, B.; Wang, W.; Guan, X.; Ouyang, Q.; Li, L.; Zhai, J.; Li, C.; Li, L.; Xia, X.; Yang, L.; Qian, H.; Yi, X.; Xu, B.; Ma, F. Molecular Landscape and Efficacy of HER2-Targeted Therapy in Patients with HER2-Mutated Metastatic Breast Cancer. Npj Breast Cancer 2020, 6 (1), 1–8. 10.1038/s41523-020-00201-9.

(73) Marusyk, A.; Janiszewska, M.; Polyak, K. Intratumor Heterogeneity: The Rosetta Stone of Therapy Resistance. Cancer Cell 2020, 37 (4), 471–484. 10.1016/j.ccell.2020.03.007.

(74) Cannavo, E.; Gerrits, B.; Marra, G.; Schlapbach, R.; Jiricny, J. Characterization of the Interactome of the Human MutL Homologues MLH1, PMS1, and PMS2. J. Biol. Chem. 2007, 282 (5), 2976–2986. 10.1074/jbc.M609989200.

(75) Pannafino, G.; Alani, E. Coordinated and Independent Roles for MLH Subunits in DNA Repair. Cells 2021, 10 (4), 948. 10.3390/cells10040948.

(76) Furman, C. M.; Wang, T.-Y.; Zhao, Q.; Yugandhar, K.; Yu, H.; Alani, E. Handcuffing Intrinsically Disordered Regions in Mlh1–Pms1 Disrupts Mismatch Repair. Nucleic Acids Res. 2021, 49 (16), 9327–9341. 10.1093/nar/gkab694.

(77) Hall, M. C.; Wang, H.; Erie, D. A.; Kunkel, T. A. High Affinity Cooperative DNA Binding by the Yeast Mlh1-Pms1 Heterodimer. J. Mol. Biol. 2001, 312 (4), 637–647. 10.1006/jmbi.2001.4958.

(78) Hombauer, H.; Campbell, C. S.; Smith, C. E.; Desai, A.; Kolodner, R. D. Visualization of Eukaryotic DNA Mismatch Repair Reveals Distinct Recognition and Repair Intermediates. Cell 2011, 147 (5), 1040–1053. 10.1016/j.cell.2011.10.025.

(79) Ortega, J.; Lee, G. S.; Gu, L.; Yang, W.; Li, G.-M. Mispair-Bound Human MutS–MutL Complex Triggers DNA Incisions and Activates Mismatch Repair. Cell Res. 2021, 31 (5), 542–553. 10.1038/s41422-021-00468-y.

(80) Witte, S. J.; Rosa, I. M.; Collingwood, B. W.; Piscitelli, J. M.; Manhart, C. M. The Mismatch Repair Endonuclease MutLα Tethers Duplex Regions of DNA Together and Relieves DNA Torsional Tension. Nucleic Acids Res. 2023, 51 (6), 2725–2739. 10.1093/nar/gkad096.

(81) Dutta, R.; Inouye, M. GHKL, an Emergent ATPase/Kinase Superfamily. Trends Biochem. Sci. 2000, 25 (1), 24–28. 10.1016/S0968-0004(99)01503-0.

(82) Wu, H.; Zeng, H.; Lam, R.; Tempel, W.; Kerr, I. D.; Min, J. Structure of the Human MLH1 N-Terminus: Implications for Predisposition to Lynch Syndrome. Acta Crystallogr. Sect. F Struct. Biol. Commun. 2015, 71 (8), 981–985. 10.1107/S2053230X15010183.

(83) Wu, X.; Platt, J. L.; Cascalho, M. Dimerization of MLH1 and PMS2 Limits Nuclear Localization of MutLα. Mol. Cell. Biol. 2003, 23 (9), 3320–3328. 10.1128/MCB.23.9.3320-3328.2003.

(84) de Barros, A. C.; Takeda, A. A. S.; Dreyer, T. R.; Velazquez-Campoy, A.; Kobe, B.; Fontes, M. R. M. DNA Mismatch Repair Proteins MLH1 and PMS2 Can Be Imported to the Nucleus by a Classical Nuclear Import Pathway. Biochimie 2018, 146, 87–96. 10.1016/j.biochi.2017.11.013.

(85) Sacho, E. J.; Kadyrov, F. A.; Modrich, P.; Kunkel, T. A.; Erie, D. A. Direct Visualization of Asymmetric Adenine-Nucleotide-Induced Conformational Changes in MutL Alpha. Mol. Cell 2008, 29 (1), 112–121. 10.1016/j.molcel.2007.10.030.

(86) Kim, Y.; Furman, C. M.; Manhart, C. M.; Alani, E.; Finkelstein, I. J. Intrinsically Disordered Regions Regulate Both Catalytic and Non-Catalytic Activities of the MutLα Mismatch Repair Complex. Nucleic Acids Res. 2019, 47 (4), 1823–1835. 10.1093/nar/gky1244.

(87) Mardenborough, Y. S. N.; Nitsenko, K.; Laffeber, C.; Duboc, C.; Sahin, E.; Quessada-Vial, A.; Winterwerp, H. H. K.; Sixma, T. K.; Kanaar, R.; Friedhoff, P.; Strick, T. R.; Lebbink, J. H. G. The Unstructured Linker Arms of MutL Enable GATC Site Incision beyond Roadblocks during Initiation of DNA Mismatch Repair. Nucleic Acids Res. 2019, 47 (22), 11667–11680. 10.1093/nar/gkz834.

(88) Schmidt, M. K.; Tollenaar, R. A. E. M.; de Kemp, S. R.; Broeks, A.; Cornelisse, C. J.; Smit, V. T. H. B. M.; Peterse, J. L.; van Leeuwen, F. E.; Van’t Veer, L. J. Breast Cancer Survival and Tumor Characteristics in Premenopausal Women Carrying the CHEK2*1100delC Germline Mutation. J. Clin. Oncol. Off. J. Am. Soc. Clin. Oncol. 2007, 25 (1), 64–69. 10.1200/JCO.2006.06.3024.

(89) Mayer, E. L.; Dueck, A. C.; Martin, M.; Rubovszky, G.; Burstein, H. J.; Bellet-Ezquerra, M.; Miller, K. D.; Zdenkowski, N.; Winer, E. P.; Pfeiler, G.; Goetz, M.; Ruiz-Borrego, M.; Anderson, D.; Nowecki, Z.; Loibl, S.; Moulder, S.; Ring, A.; Fitzal, F.; Traina, T.; Chan, A.; Rugo, H. S.; Lemieux, J.; Henao, F.; Lyss, A.; Novoa, S. A.; Wolff, A. C.; Vetter, M.; Egle, D.; Morris, P. G.; Mamounas, E. P.; Gil-Gil, M. J.; Prat, A.; Fohler, H.; Filho, O. M.; Schwarz, M.; DuFrane, C.; Fumagalli, D.; Theall, K. P.; Lu, D. R.; Bartlett, C. H.; Koehler, M.; Fesl, C.; DeMichele, A.; Gnant, M. Palbociclib with Adjuvant Endocrine Therapy in Early Breast Cancer (PALLAS): Interim Analysis of a Multicentre, Open-Label, Randomised, Phase 3 Study. Lancet Oncol. 2021, 22 (2), 212–222. 10.1016/S1470-2045(20)30642-2.

(90) Loibl, S.; Marmé, F.; Martin, M.; Untch, M.; Bonnefoi, H.; Kim, S.-B.; Bear, H.; McCarthy, N.; Melé Olivé, M.; Gelmon, K.; García-Sáenz, J.; Kelly, C. M.; Reimer, T.; Toi, M.; Rugo, H. S.; Denkert, C.; Gnant, M.; Makris, A.; Koehler, M.; Huang-Bartelett, C.; Lechuga Frean, M. J.; Colleoni, M.; Werutsky, G.; Seiler, S.; Burchardi, N.; Nekljudova, V.; von Minckwitz, G. Palbociclib for Residual High-Risk Invasive HR-Positive and HER2-Negative Early Breast Cancer—The Penelope-B Trial. J. Clin. Oncol. 2021, 39 (14), 1518–1530. 10.1200/JCO.20.03639.

(91) Johnston, S. R. D.; Toi, M.; O’Shaughnessy, J.; Rastogi, P.; Campone, M.; Neven, P.; Huang, C.-S.; Huober, J.; Jaliffe, G. G.; Cicin, I.; Tolaney, S. M.; Goetz, M. P.; Rugo, H. S.; Senkus, E.; Testa, L.; Mastro, L. D.; Shimizu, C.; Wei, R.; Shahir, A.; Munoz, M.; Antonio, B. S.; André, V.; Harbeck, N.; Martin, M. Abemaciclib plus Endocrine Therapy for Hormone Receptor-Positive, HER2-Negative, Node-Positive, High-Risk Early Breast Cancer (MonarchE): Results from a Preplanned Interim Analysis of a Randomised, Open-Label, Phase 3 Trial. Lancet Oncol. 2023, 24 (1), 77–90. 10.1016/S1470-2045(22)00694-5.

(92) Slamon, D. J.; Stroyakovskiy, D.; Yardley, D. A.; Huang, C.-S.; Fasching, P. A.; Crown, J.; Bardia, A.; Chia, S.; Im, S.-A.; Martin, M.; Loi, S.; Xu, B.; Hurvitz, S. A.; Barrios, C.; Untch, M.; Moroose, R. L.; Visco, F.; Fresco, R.; Taran, T.; Hortobagyi, G. N. Ribociclib and Endocrine Therapy as Adjuvant Treatment in Patients with HR+/HER2- Early Breast Cancer: Primary Results from the Phase III NATALEE Trial. J. Clin. Oncol. 2023, 41 (17_suppl), LBA500–LBA500. 10.1200/JCO.2023.41.17_suppl.LBA500.

(93) Asghar, U. S.; Kanani, R.; Roylance, R.; Mittnacht, S. Systematic Review of Molecular Biomarkers Predictive of Resistance to CDK4/6 Inhibition in Metastatic Breast Cancer. *JCO Precis*. Oncol. 2022, No. 6, e2100002. 10.1200/PO.21.00002.

(94) Bui, T. B. V.; Burgering, B. M. T.; Goga, A.; Rugo, H. S.; van ‘t Veer, L. J. Biomarkers for Cyclin-Dependent Kinase 4/6 Inhibitors in the Treatment of Hormone Receptor-Positive/Human Epidermal Growth Factor Receptor 2-Negative Advanced/Metastatic Breast Cancer: Translation to Clinical Practice. JCO Precis. Oncol. 2022, No. 6, e2100473. 10.1200/PO.21.00473.

(95) Selli, C.; Dixon, J. M.; Sims, A. H. Accurate Prediction of Response to Endocrine Therapy in Breast Cancer Patients: Current and Future Biomarkers. Breast Cancer Res. 2016, 18 (1), 118. 10.1186/s13058-016-0779-0.

(96) Portman, N.; Alexandrou, S.; Carson, E.; Wang, S.; Lim, E.; Caldon, C. E. Overcoming CDK4/6 Inhibitor Resistance in ER-Positive Breast Cancer. Endocr. Relat. Cancer 2019, 26 (1), R15–R30. 10.1530/ERC-18-0317.

(97) Staff, T. A. P. Companion Diagnostic to Identify Patients With Endometrial Cancer Eligible for Pembrolizumab Therapy Approved by the FDA - The ASCO Post. https://ascopost.com/issues/september-10-2022/companion-diagnostic-to-identify-patients-with-endometrial-cancer-eligible-for-pembrolizumab-therapy-approved-by-the-fda/ (accessed 2024-02-12).

(98) Vikas, P.; Messersmith, H.; Compton, C.; Sholl, L.; Broaddus, R. R.; Davis, A.; Estevez-Diz, M.; Garje, R.; Konstantinopoulos, P. A.; Leiser, A.; Mills, A. M.; Norquist, B.; Overman, M. J.; Sohal, D.; Turkington, R. C.; Johnson, T. Mismatch Repair and Microsatellite Instability Testing for Immune Checkpoint Inhibitor Therapy: ASCO Endorsement of College of American Pathologists Guideline. J. Clin. Oncol. 2023, 41 (10), 1943–1948. 10.1200/JCO.22.02462.

