## Supplementary files for "Aberrant cytoplasmic localization of MLH1 characterizes a sub-clonal breast cancer cell population that seeds recurrence"

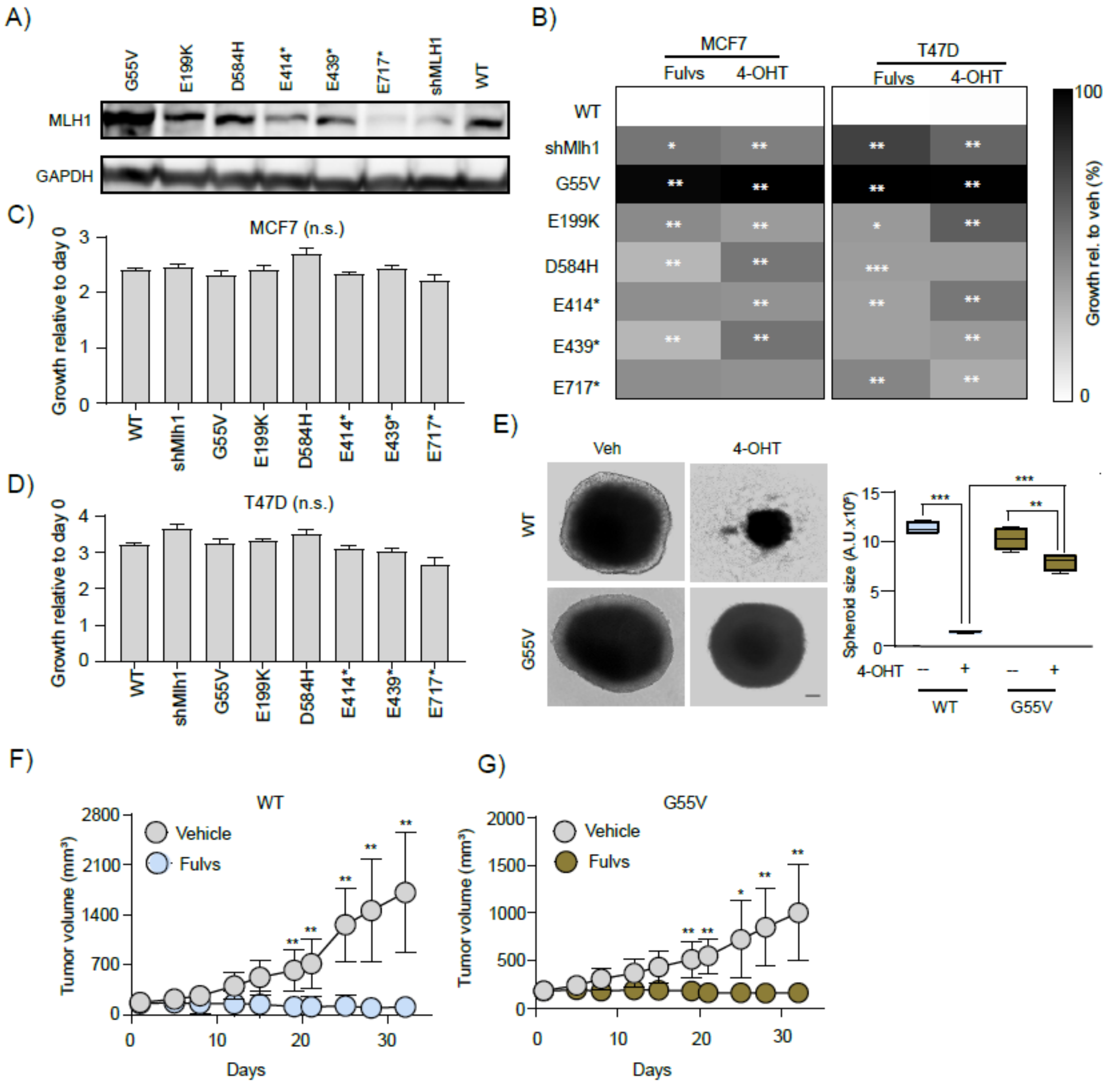


**Figure S1: Some *MLH1* mutations induce endocrine therapy resistance in breast cancer cell lines.** (A) Western blot confirming stable expression of MLH1 mutant protein in MCF7 cells stably expressing shRNA against *MLH1*. (B) Heatmap showing viability of MCF7 and T47D cells stably expressing either mutant or wild type constructs after fulvestrant (100nM) or 4-OHT (100nM) treatment. (C-D) Bar graph showing basal level growth of MCF7 and T47D cells stably expressing *MLH1* mutant or WT constructs. Data expressed as fold change relative to day 0 reading. Supports data presented in Fig 1A-D using transiently transfected cells. (E) Representative images of 3D growth assay of T47D$^{G55V}$ and T47D$^{WT}$ spheroids after 4-(OHT) treatment. Accompanying box plot with quantification. Supports data representing response to fulvestrant presented in Fig 1E-F. (F-G) Dot plot representing the average volumes of tumors arising from MCF7$^{WT}$ (F) and MCF7$^{G55V}$ (G) cells xenografted into nude mice with or without fulvestrant treatment. Supports normalized data presented in Fig 1I. Student's t-test determined all p-values. *** represents p≤0.0001, ** p≤0.01 and * p≤0.05. For all graphs, error bars describe standard deviation.

Scale bar = 50μm. Abbreviation Vehicle (Veh), Fulvestrant (Fulvs), Tamoxifen (4-OHT), Wild type (WT), not significant (n.s.).

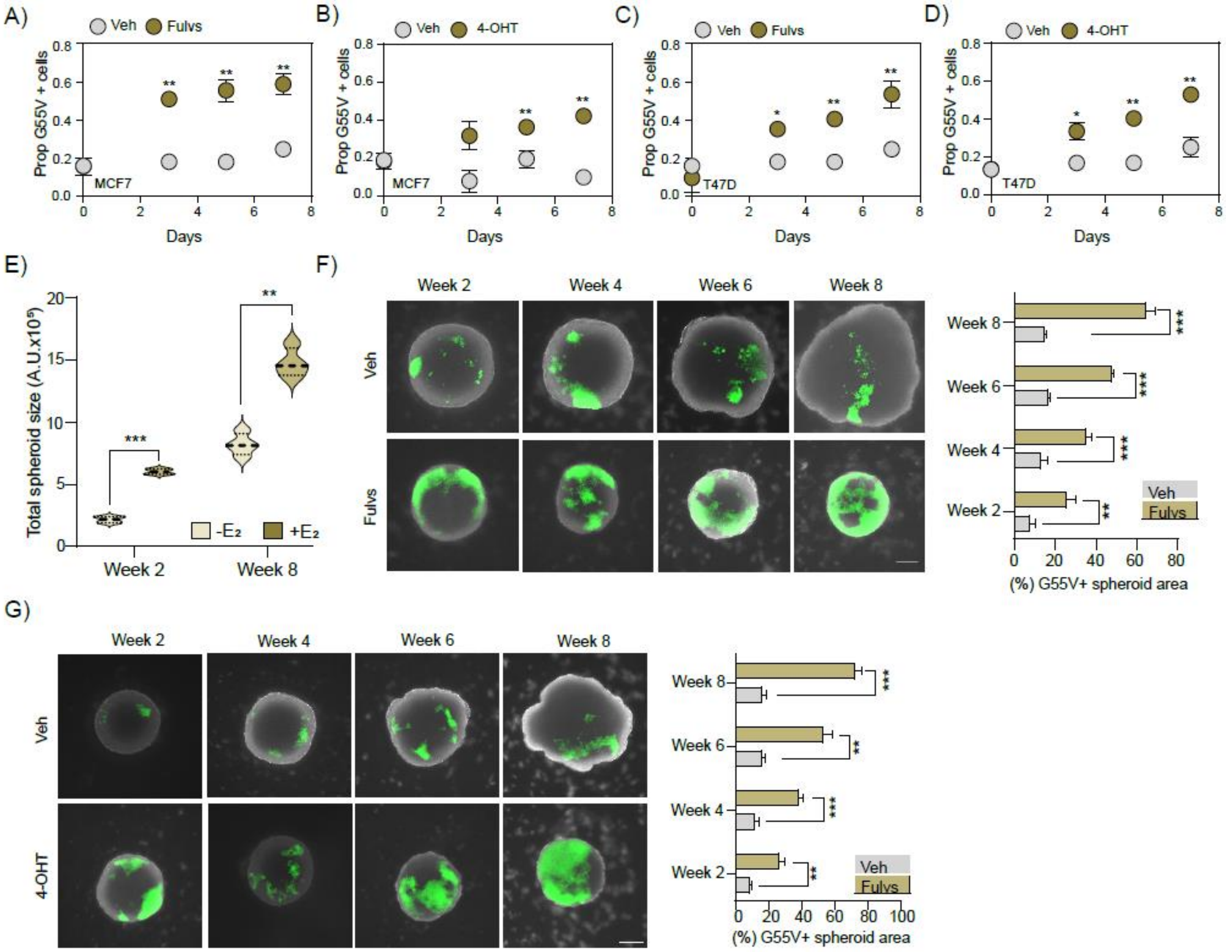


**Figure S2: *MLH1$^{G55V}$* cells persist in subclonal fractions after endocrine therapy**. (A-D) Dot plots showing increase in GFP+ MCF7$^{G55V}$ (A-B) and T47D$^{G55V}$ (C-D) cells after fulvestrant (A,C) or tamoxifen (B, D) treatment as indicated. Supports estrogen deprivation data presented in Fig 3C-D. (E) Violin plot showing change in total spheroid size of T47D cells with or without estrogen supplementation. Supports data presented in Fig 3E. (F-G) Representative images of 3D spheroids comprised of T47D$^{G55V}$ :T47D$^{WT}$ cells (1:9) grown with or without fulvestrant (F) and tamoxifen (G) for 8 weeks with accompanying quantification of GFP+ area as bar graphs. Supports data presented in Fig 3F. Green = GFP. Student's t-test determined p-values. *** represents $p \leq 0.0001$, ** $p \leq 0.01$ and * $p \leq 0.05$. For all graphs, error bars describe standard deviation. Scale bar = 20μm. Abbreviations, Vehicle (Veh), Wild type (WT), tamoxifen (4-OHT), not significant (n.s.).

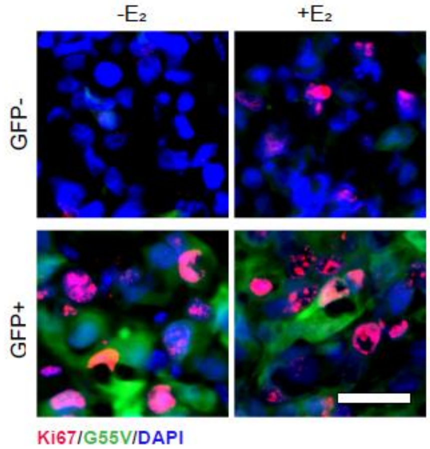


**Figure S3: MLH1$^{G55V}$ cells persist sub-clonally after endocrine deprivation *in vivo*.** Representative photomicrographs for Ki67+/GFP+ (G55V) and Ki67+/GFP- (WT) cells in both +$E_2$ and -$E_2$ conditions. Supports data presented in Fig 3I. Scale bar = 20μm. Abbreviations, ($E_2$) beta-estradiol.

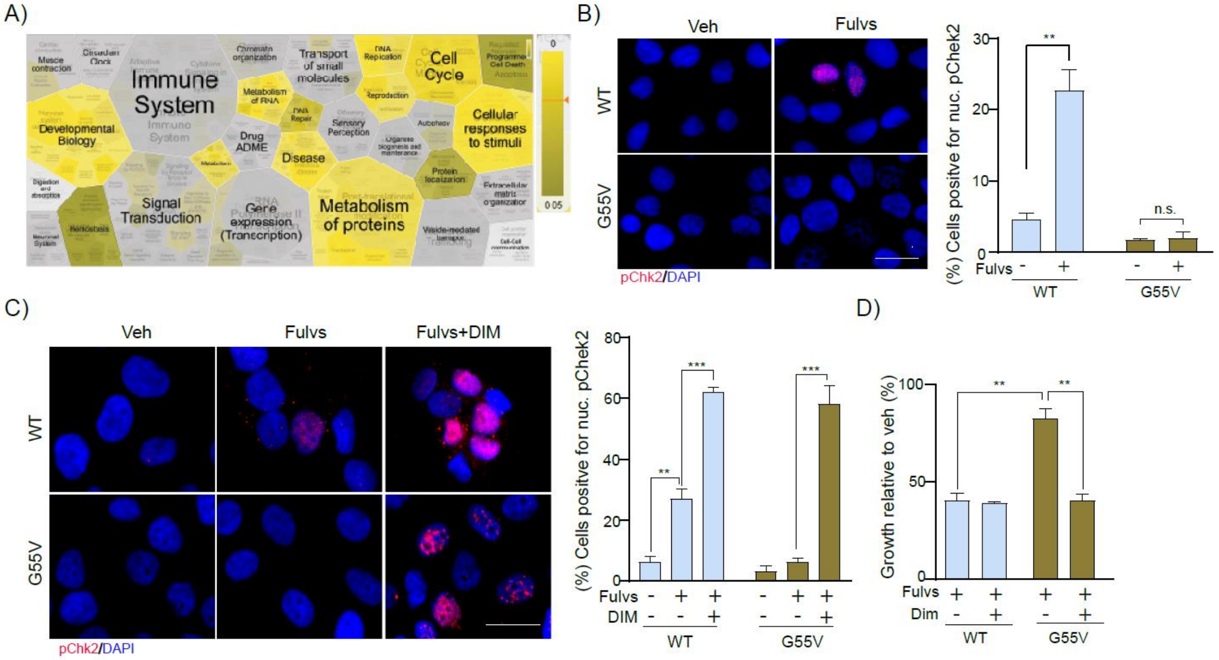


**Figure S4: *MLH1$^{G55V}$* prevents Chk2 activation.** (A) Reactome analysis of RNAseq data from MCF7 cells after 1μM fulvestrant treatment for pathways that are enriched for up- or down-regulation in response to fulvestrant specifically in G55V cells but not in WT-, shMLH1- or *MLH1$^{E717*}$*-bearing cells. Supports data presented in Fig 4A. (B) Representative immunofluorescence pictures showing pChk2 subcellular localization in T47D cells stably expressing *MLH1$^{G55V}$* or *MLH1$^{WT}$* with or without treatment with fulvestrant 1μM for 36h. (C) Representative immunofluorescence pictures showing pChk2 expression in T47D$^{G55V}$ and T47D$^{WT}$ cells stably after 1μM fulvestrant treatment alone or combinatorial fulvestrant (100nM) + DIM (200μM) treatment for 36h. Accompanying quantification presented as bar graphs. (D) Bar graphs showing growth of T47D$^{G55V}$ and T47D$^{WT}$ cells after either 1μM fulvestrant treatment alone or combinatorial fulvestrant (100nM) + DIM (200μM) treatment for 36h. Growth is normalized to vehicle for each type and expressed as percent inhibition. Student's t-test was performed to determine all p-values. Panels B-D support data from MCF7 cells presented in Fig 4B-C. *** represents p≤0.0001, ** p≤0.01 and * p≤0.05. For all graphs, error bars describe standard deviation. Scale bar = 20μm. Abbreviations, Vehicle (Veh), Fulvestrant (Fulvs), 3,3′-Di-indolylmethane (DIM).

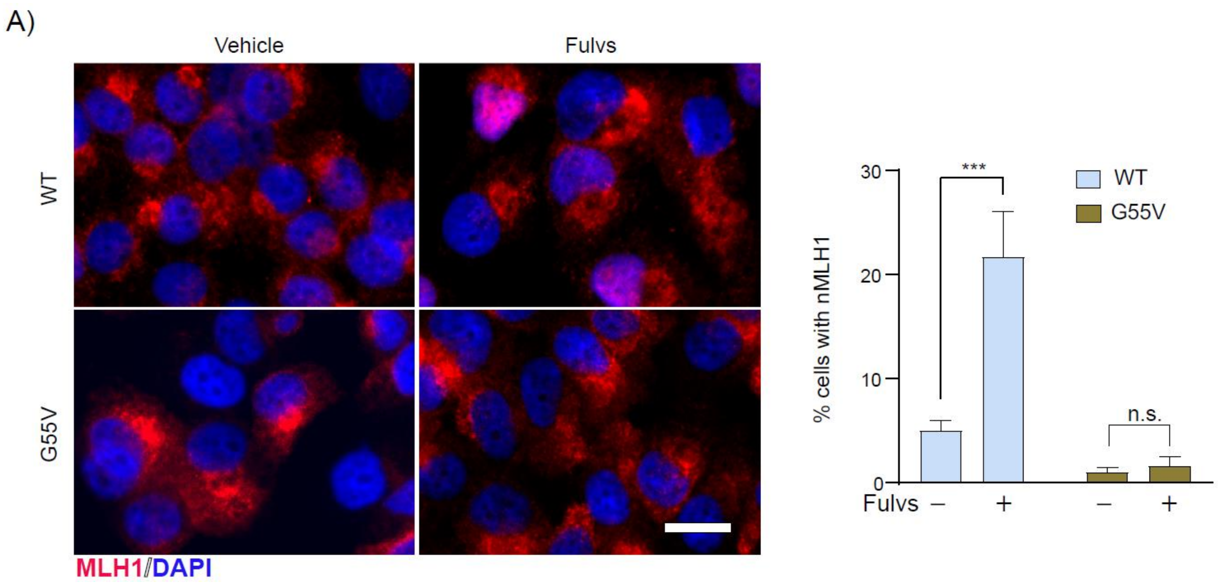


**Figure S5: G55V mutation alters subcellular localization of MLH1 protein in T47D cells.** Representative immunofluorescence pictures showing subcellular expression of MLH1 in T47D$^{G55V}$ or T47D$^{WT}$ cells after 1μM fulvestrant treatment for 36h. Accompanying bar graph shows quantification. Student's t-test determined all p-values. *** represents $p \leq 0.0001$, ** $p \leq 0.01$ and * $p \leq 0.05$. For graph, error bars describe standard deviation. Scale bar = 50μm. Abbreviations, Vehicle (Veh), Wild type (WT), Fulvestrant (Fulvs), not significant (n.s.). Supports data in MCF7 cells presented in Fig 5.

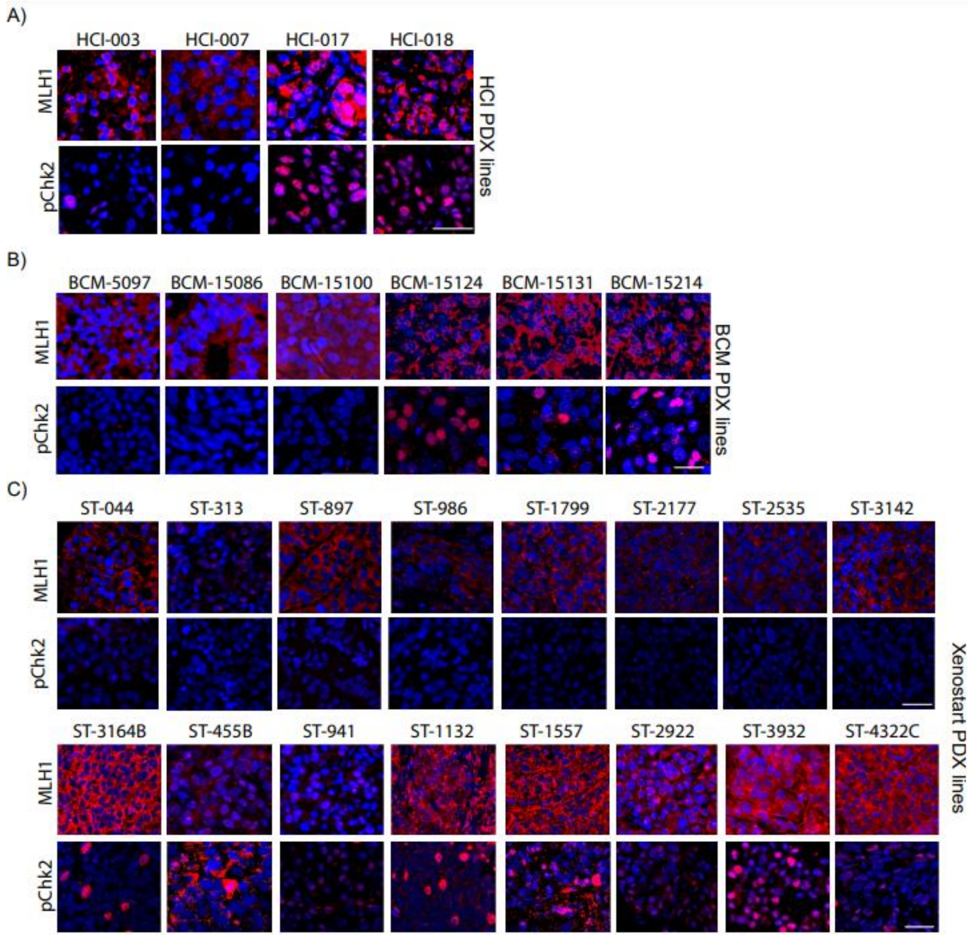


**Figure S6: Cytoplasmic MLH1 reduces Chk2 activation in multiple cohorts of ER+/HER2- PDX.** Representative immunofluorescence images of MLH1 and pChk2 sub-cellular localization in HCI (A), BCM (B) and Xenostart PDX lines (C). Scale bars = 20µm. Quantification of immunofluorescence is presented in Fig 6. Abbreviations, HCI (Huntsman Cancer Institute), BCM (Baylor College of Medicine).

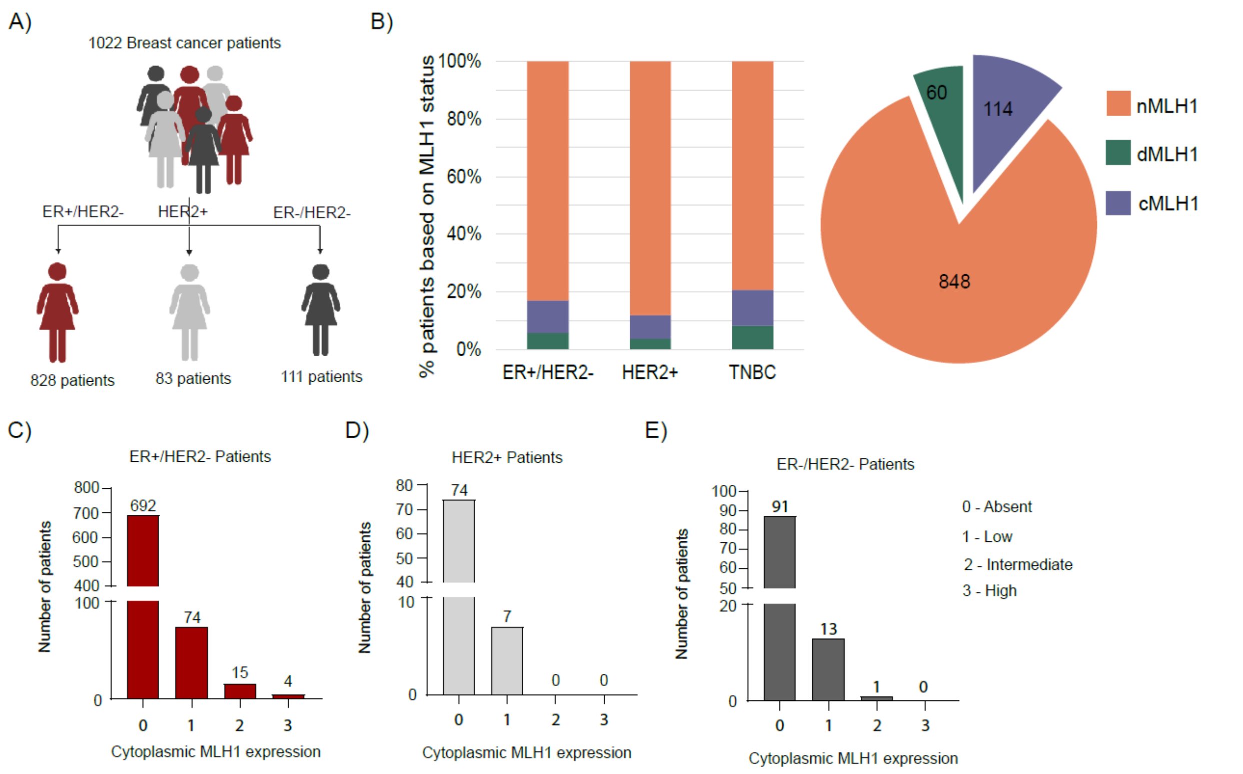


**Figure S7: Subcellular expression of MLH1 in ER+/HER2- patient biopsies.** (A) Description of patient tumor dataset used in Fig 5F. (B) Stacked column graph and pie chart indicating proportion of patient tumors with MLH1 staining characterized as cytoplasmic (1-3 score for cytoplasmic MLH1 staining), nuclear (0 score for cytoplasmic MLH1), or deficient (no detectable MLH1 positivity) overall (pie chart) and by breast cancer type (stacked column). Representative images of cytoplasmic and nuclear MLH1 are presented in Fig 5F. (C-E) Cytoplasmic MLH1 expression graded on a scale of 0 to 3 with (0) Absent, (1) Low, (2) Intermediate, (3) High for patient tumors of delineated subtypes.

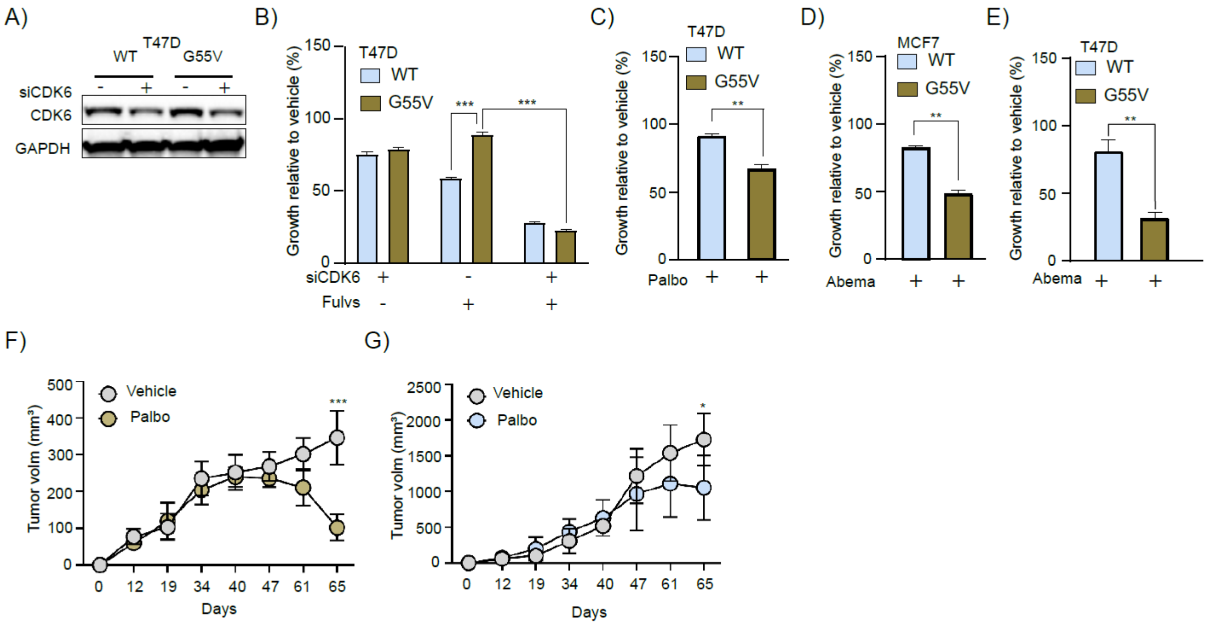


**Figure S8: Cytoplasmic MLH1 promotes sensitivity to CDK4/6 inhibition.** (A) Western blot confirming knockdown of *CDK6* in T47D$^{WT}$ and T47D$^{G55V}$ cells. Supports data for MCF7 cells presented in Fig 6C. (B-E) Bar graphs showing growth of T47D$^{WT}$ and T47D$^{G55V}$ cells (B-C, E) and MCF7$^{WT}$ and MCF7$^{G55V}$ cells (D) after *CDK6* knockdown alone or with fulvestrant treatment (B) or after treatment with CDK4/6 inhibitors palbociclib (C) or abemaciclib (D-E). Supports data for MCF7 cells presented in Fig 6D-E. (F-G) Average volume of tumors from MCF7$^{G55V}$ (F) and MCF7$^{WT}$ (G) cells xenografted into nude mice with or without palbociclib treatment over time. Supports normalized data presented in Fig 6G. For graphs errors bars describe standard deviation. Student's t-test determined all p-values. *** represents $p \leq 0.0001$, ** $p \leq 0.01$ and *$p \leq 0.05$.
